# Cwl0971, a novel peptidoglycan hydrolase, plays pleiotropic roles in *Clostridioides difficile* R20291

**DOI:** 10.1101/2020.11.29.402867

**Authors:** Duolong Zhu, Hiran Malinda Lamabadu Warnakulasuriya Patabendige, Brooke Rene Tomlinson, Shaohui Wang, Syed Hussain, Domenica Flores, Yongqun He, Lindsey N Shaw, Xingmin Sun

## Abstract

*Clostridioides difficile* is a Gram-positive, spore-forming, toxin-producing anaerobe that can cause nosocomial antibiotic-associated intestinal disease. Although the production of toxin A (TcdA) and toxin B (TcdB) contribute to the main pathogenesis of *C. difficile*, the mechanism of TcdA and TcdB release from cell remains unclear. In this study, we identified and characterized a new cell wall hydrolase Cwl0971 (*CDR20291_0971*) from *C. difficile* R20291, which is involved in bacterial autolysis. The gene 0971 deletion mutant (R20291Δ0971) generated with CRISPR-AsCpfI exhibited significantly delayed cell autolysis and increased cell viability compared to R20291, and the purified Cwl0971 exhibited hydrolase activity for *Bacillus subtilis* cell wall. Meanwhile, 0971 gene deletion impaired TcdA and TcdB release due to the decreased cell autolysis in the stationary / late phase of cell growth. Moreover, sporulation of the mutant strain decreased significantly compared to the wild type strain. *In vivo*, the defect of Cwl0971 decreased fitness over the parent strain in a mouse infection model. Collectively, Cwl0971 is involved in cell wall lysis and cell viability, which affects toxin release, sporulation, germination, and pathogenicity of R20291, indicating that Cwl0971 could be an attractive target for *C. difficile* infection therapeutics and prophylactics.

## Introduction

*Clostridioides difficile* is a Gram-positive, spore-forming, toxin-producing, anaerobic bacterium that has established itself as a leading cause of nosocomial antibiotic-associated diarrhea in the developed countries (Sebaihia et al., 2006). *C. difficile* infection (CDI) can result in a spectrum of symptoms, ranging from mild diarrhea to pseudomembranous colitis and potential death (Lessa et al., 2012). Recently, morbidity and mortality rates of CDI have been increasing steadily, causing over 500,000 infections per year in the United States alone with an estimated cost of $1-3 billion (Dubberke and Olsen, 2012; Lessa et al., 2015). *C. difficile* has many virulence factors, such as toxins, adhesins, flagella, and proteases (Borriello et al., 1990; Janoir et al., 2007; Deneve et al., 2009; Stevenson et al., 2015; Janoir, 2016; Kevorkian et al., 2016; Pechine et al., 2016). Among them, two large potent exotoxins, toxin A (TcdA) and toxin B (TcdB) are the most well-studied virulence factors (Kuehne et al., 2010). These toxins can disrupt the actin cytoskeleton of intestinal epithelial cells and inate immune cells through glucosylation of the Rho family of GTPases to induce mucosal inflammation and the symptoms associated with CDI (Voth and Ballard, 2005).

The toxin encoding genes *tcdA* and *tcdB* are located in a 19.6 kb pathogenicity locus (PaLoc), which also contains three additional genes, *tcdR, tcdC*, and *tcdE* (Braun et al., 1996; Mani and Dupuy, 2001). The gene *tcdR* encodes an RNA polymerase sigma factor that positively regulates the expression of both toxin genes and its own gene (Mani et al., 2002). TcdC is a putative antagonist of TcdR that negatively regulates TcdR-containing RNA polymerase holoenzyme (Dupuy et al., 2008). While several other previous studies have showed that TcdC might have a moderate role in regulating toxin expression, it is not a major determinant of the hypervirulence of *C. difficile* (Murray et al., 2009; Bakker et al., 2012; Martin-Verstraete et al., 2016). Recently, Paiva et al. (Paiva et al., 2020) identified that the C-terminal domain of TcdC is exposed on the bacterial cell surface, which is not compatible with the direct binding of intracellular protein targets, indicating another regulation mechanism of TcdC that is different from anti-sigma factors. TcdA and TcdB belong to the large clostridial glucosylating toxin family, which are released without signal peptide (Popoff and Bouvet, 2009). TcdE, the holin-like protein, has been reported as being involved in toxin release (Govind and Dupuy, 2012; Govind et al., 2015). However, the results of TcdE studies from different groups are controversial. Olling et al. (Olling et al., 2012) reported that toxin release from *C. dificile* 630Δ*erm* is not affected by *tcdE* gene mutation and indicated that transfer of *tcdE* gene into other microorganisms could lead to spontaneous lysis of the recipient. Recently, Wydau-Dematteis et al. (Wydau-Dematteis et al., 2018) characterized Cwp19, a novel lytic transglycosylase, which can contribute to toxin release through bacteriolysis. In our previous study, we identified a novel peptidoglycan cross-linking enzyme Cwp22, which is involved in *C. difficile* cell wall integrity and can affect toxin release (Zhu et al., 2019). A link between toxin release and cell wall hydrolase-induced bacteriolysis needs further investigation to uncover the toxin secretion mechanism.

Peptidoglycan (PG), an essential cell wall biopolymer, is a primary cell wall constituent of Gram-positive bacteria formed by glycan strands of β-1,4-linked-N-acetylmuramic acid (NAM) and N-acetylglucosamine (NAG), cross-linked by short peptides attached to NAM (Layec et al., 2008). PG is a dynamic macromolecule and the constant equilibrium of PG in the cell is strictly controlled to enable bacterial growth and differentiation (Layec et al., 2008). Peptidoglycan hydrolases (PGHs, including bacteria autolysins) contribute to PG plasticity for maintaining cell wall shape through hydrolyzing PG bonds (Layec et al., 2008). It is reasonably clear that PGHs play the key roles in the processes of (I) regulation of cell wall growth, (II) turnover of peptidoglycan, (III) separation of daughter cells, (□) liberation of spores from mother cells, (□) germination of spores to form vegetative cells, and (□) bacteria autolysis (Layec et al., 2008; Vollmer et al., 2008). Regulation of PGHs in these processes could affect the pathogenic properties of some bacteria to their hosts. According to the substrate specificity and the resulting cleavage products, PGHs can be defined into 5 classes as N-acetylmuramoyl-L-alanine amidases (also named as amidases), N-acetylglucosaminidases, N-acetylmuramidases (lysozyme and transglycosylases included), carboxypeptidases, and endopeptidases (Layec et al., 2008; Vollmer et al., 2008). Among them, endopeptidases can be sub-classed as D,D-, L,D-, and D,L-endopeptidase referring to the stereochemistry of the cleaved amino acid residues. In Gram-positive bacteria, Amidases AtlA and SleI can affect bacterial biofilm formation, autolysis, and virulence in *Staphylococcus aureus* (Bose et al., 2012; Vermassen et al., 2019); lytic enzyme CbpD, LytA, LytB, and LytC can mediate lysis of non-competent target cells and cell separation in *Streptococcus pneumoniae* (Lopez et al., 2000; Eldholm et al., 2009); N-acetylglucosaminidase (p60, FlaA) and N-acetylmuramoyl-L-alanine amidases (Ami, IspC) can affect cell separation, adhesion, virulence, and autolysis in *Listeria monocytogenes* (Wuenscher et al., 1993; Milohanic et al., 2001; Popowska and Markiewicz, 2004; Wang and Lin, 2008). Carboxypeptidase Pgp1 and Pgp2 in Gram-negative strain *Campylobacter jejuni* could reduce pathogenicity through impairing bacteria motility, biofilm formation, and bacteria colonization (Frirdich et al., 2012; Frirdich et al., 2014).

PGHs have been studied well in many pathogens, while the characterization and physiological functions of PGHs in *C. difficile* remains to be further explored. In total, 37 PGHs belonging to each of the five hydrolase classes were predicted in *C. difficile* 630, among them are 13 endopeptidases, 10 N-acetylmuramoyl-L-alanine amidases, 8 carboxypeptidases, 3 N-acetylglucosaminidases, and 3 N-acetylmuramidases (Layec et al., 2008). Acd (N-acetylglucosaminidases, autolysin) which can hydrolyze peptidoglycan bonds between N-acetylglucosamine and N-acetylmuramic acid and CwlT hydrolase which contains a novel bacterial lysozyme domain and an NlpC/P60 D,L-endopeptidase domain were firstly identified in *C. difficile* 630, while the physiological functions of Acd and CwlT are still not clear (Dhalluin et al., 2005; Xu et al., 2014). Besides, SleC, a N-acetylglucosaminidases which is essential for *C. difficile* spore germination, and SpoIIQ, an endopeptidase that is required for *C. difficile* spore formation have also been identified (Gutelius et al., 2014; Fimlaid et al., 2015). Recently, Cwp19 and Cwp22 which can affect toxin release through bacteriolysis and cell wall integrity change were also reported in *C. difficile* (Wydau-Dematteis et al., 2018; Zhu et al., 2019).

Although several PGHs have been reported in *C. difficile*, more putative PGHs and their physiological functions need to be uncovered. In this study, we applied the Vaxign reverse vaccinology tool and characterized a new peptidoglycan hydrolase (*CDR20291_0971*, referred as Cwl0971) *in vitro* and *in vivo*. Our data showed that Cwl0971 is involved in cell wall lysis and cell viability, which affects toxin release, sporulation, germination, and pathogenicity of R20291, indicating that Cwl0971 could be an attractive target for *C. difficile* infection therapeutics and prophylactics.

## Results

### Bioinformatic identification and analysis of putative cell wall hydrolase 0971 (Cwl0971)

Vaxign is a web-based reverse vaccinology tool that uses comparative genomic sequence analysis to predict vaccine candidates based on different criteria such as cellular localization and adhesion probability (He et al., 2010). Using R20291 as the seed strain, Vaxign analysis predicted 31 *C. difficile* proteins to be cell wall-bound, likely to be adhesins and hydrolases, and conserved in the other 12 *C. difficile* genomes (file:///C:/Users/nk3ml/%E6%A1%8C%E9%9D%A2/C.%20difficile%20cell%20wall%20protein-%20vaxgin_prediction-31.htm). Among these proteins is YP_003217470.1, a putative cell wall hydrolase of 427 amino acids with a predicted molecular weight of 44.82 kDa and a pI of 9.30. This new putative cell wall hydrolase (Cwl0971) classed into C40 family peptidase is encoded by the *CDR20291_ 0971* gene (referred as 0971, NCBI Entrez Gene ID of 8468749) in strain R20291.

Based on the conserved domain analysis, Cwl0971 has a putative 27-amino acid signal sequence and two domains (Fig. 1A). The putative C-terminal catalytic domain belongs to the Spr (COG0791) and NlpC/P60 (pfam00877) superfamilies with endopeptidase activity. The N-terminal 3 bacterial SH3_3 domains belong to the YgiM superfamily (COG3103) and are predicted to act as the targeting domains involved in bacterial cell wall recognition and binding (Xu et al., 2015).

**Fig. 1.**
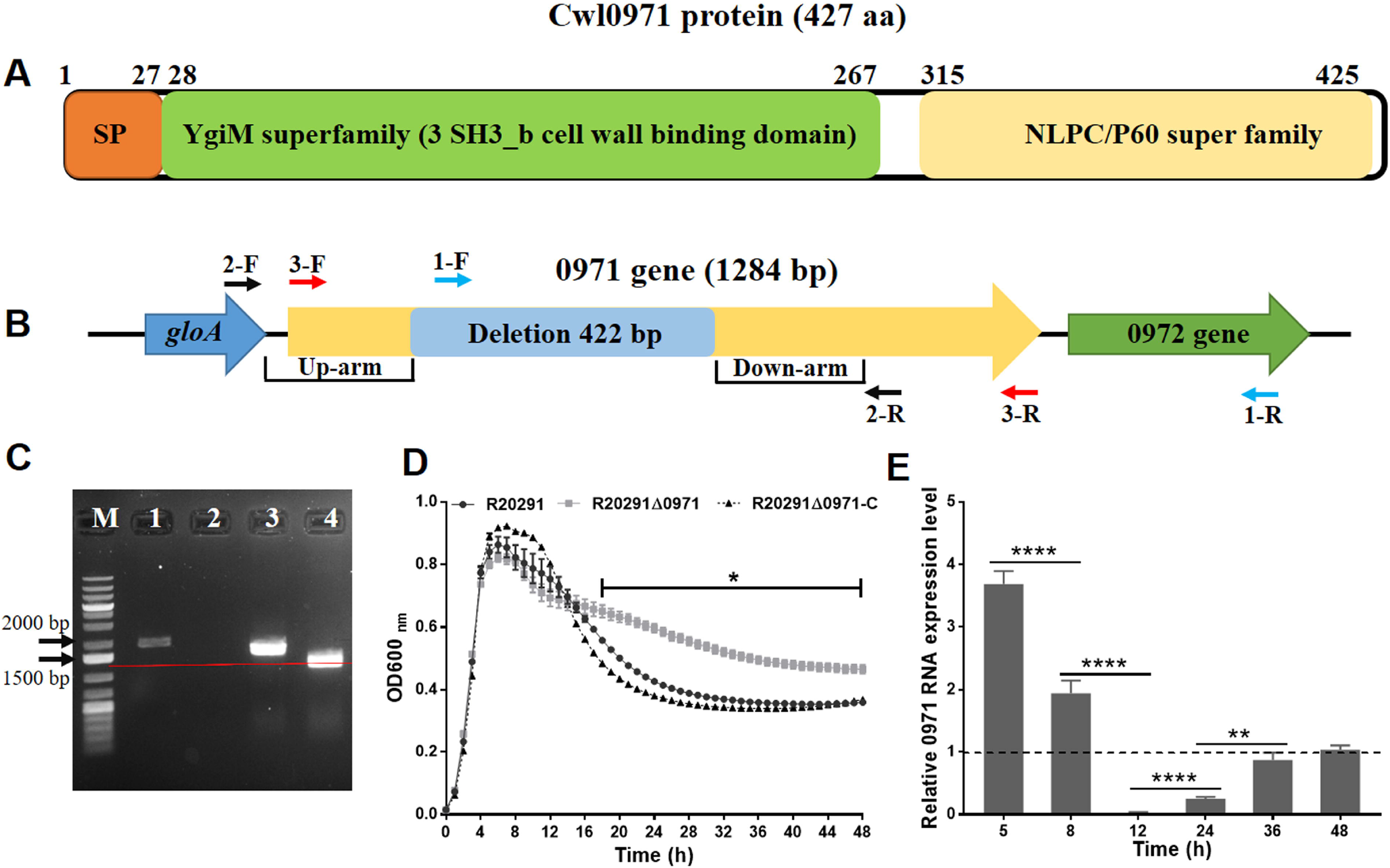
Construction of 0971 gene deletion mutant and analysis of growth profile and 0971 gene transcription. A. Schematic representation of Cwl0971 structure. Cwl0971 contains a 27 amino acid signal peptide, and two main domains that are N-terminal 3 repeat bacterial SH3_3 domain (YgiM superfamily) and C-terminal NLPC/P60 catalytic domain. B. Schematic representation of 0971 gene deletion and amplification in R20291. 1-F/R and 2-F/R were used as gene deletion verification primers; Primer 3-F/R were used for the full 0971 gene amplification. C. Verification of 0971 gene deletion in R20291. M: 1 kb DNA ladder; 1: R20291 genome PCR test with primer 1-F/R; 2: R20291Δ0971 genome PCR test with primer 1-F/R; 3: R20291 genome PCR test with primer 2-F/R; 4: R20291Δ0971 genome PCR test with primer 2-F/R. D. Growth profile of *C. difficile* strains in BHIS for 48 h. E. Transcription of 0971 gene during R20291 growth. Fold change of 0971 gene transcription at different time points was normalized to 48 h of 0971 gene transcription level. Experiments were independently repeated thrice. Bars stand for mean ± SEM. One-way analysis of variance (ANOVA) with post-hoc Tukey test was used for statistical significance, statistically significant outcomes are indicted (*\*P* < 0.05, *\*\*P* < 0.01, *\*\*\*P* < 0.001).

### Construction of 0971 gene deletion mutant and analysis of growth profile and 0971 gene transcription

To analyze the 0971 gene function in R20291, CRISPR-AsCpfI based plasmid pDL1 (pMTL82151-Ptet-AscpfI) was constructed for target gene deletion. Two sgRNAs targeting the 0971 gene in different loci were designed. A fragment of 422 bp of the 0971 gene was deleted with the pDL1-0971 plasmid, resulting in R20291Δ0971 mutant (Fig. 1 B). The correct deletion was verified with check primers 1-F/R and 2-F/R by PCR (Fig. 1 C). The 0971 gene complementation strain R20291Δ0971-pMTL84153/0971 (referred as R20291Δ0971-C), and the control strain R20291-pMTL84153 and R20291Δ0971-pMTL84153 were constructed for 0971 gene function analysis.

The effect of the 0971 gene deletion on R20291 growth was evaluated in BHIS media. As there was no significant difference in bacteria growth of R20291 VS R20291-pMTL84153 and R20291Δ0971 VS R20291Δ0971-pMTL84153 (data not shown), we only showed the growth curve of R20291, R20291Δ0971, and R20291Δ0971-C in the figure. R20291Δ0971 displayed significantly decreased autolysis than R20291 and R20291Δ0971-C starting from 16 h of post incubation (Fig 1D). The 0971 gene transcription during *C. difficile* growth peaked at 5 h of bacterial growth, followed by a reduction, and reached the lowest expression level at 12 h of bacterial growth. Interestingly, the 0971 gene expression increased again after 12 h and reached the stable expression level at 48 h, though the transcription level was still lower than that of 5 h (Fig 1E). Our RT-qPCR data indicated the potential multiple roles of the 0971 gene in different stages of *C. difficile* R20291 growth.

### Detection of Cwl0971 hydrolytic activity

NlpC/P60 superfamily protein usually has endopeptidase activity which can hydrolyze the peptidoglycan to remodel the bacterial cell wall (Anantharaman and Aravind, 2003). Cwl0971 NlpC/P60 domain structure was superimposed well with *E. coli* Spr protein which acts as *γ*-D-glutamyl-L-diaminopimelic acid endopeptidase through Ipred4 software prediction (http://www.compbio.dundee.ac.uk/jpred/) despite the low level of protein sequence identity (36% identity) between Cwl0971 and Spr (Aramini et al., 2008). The peptidoglycan layer is the main content of *C. difficile* cell wall and spore cortex (Fig. 2A). To analyze the putative hydrolytic activity of Cwl0971, we overexpressed Cwl0971 as an N-terminal His-tagged fusion protein (without signal peptide) and purified it with Ni^2+^ affinity chromatography column for zymogram analysis with *Bacillus subtilis* cell wall as substrate. As shown in Fig. 2B, reaction zones on the zymogram were detected suggesting that Cwl0971 has a detectable hydrolytic activity for the *B. subtilis* cell wall. The correct protein band was also evidenced via Western blotting (Fig. 2C).

**Fig. 2.**
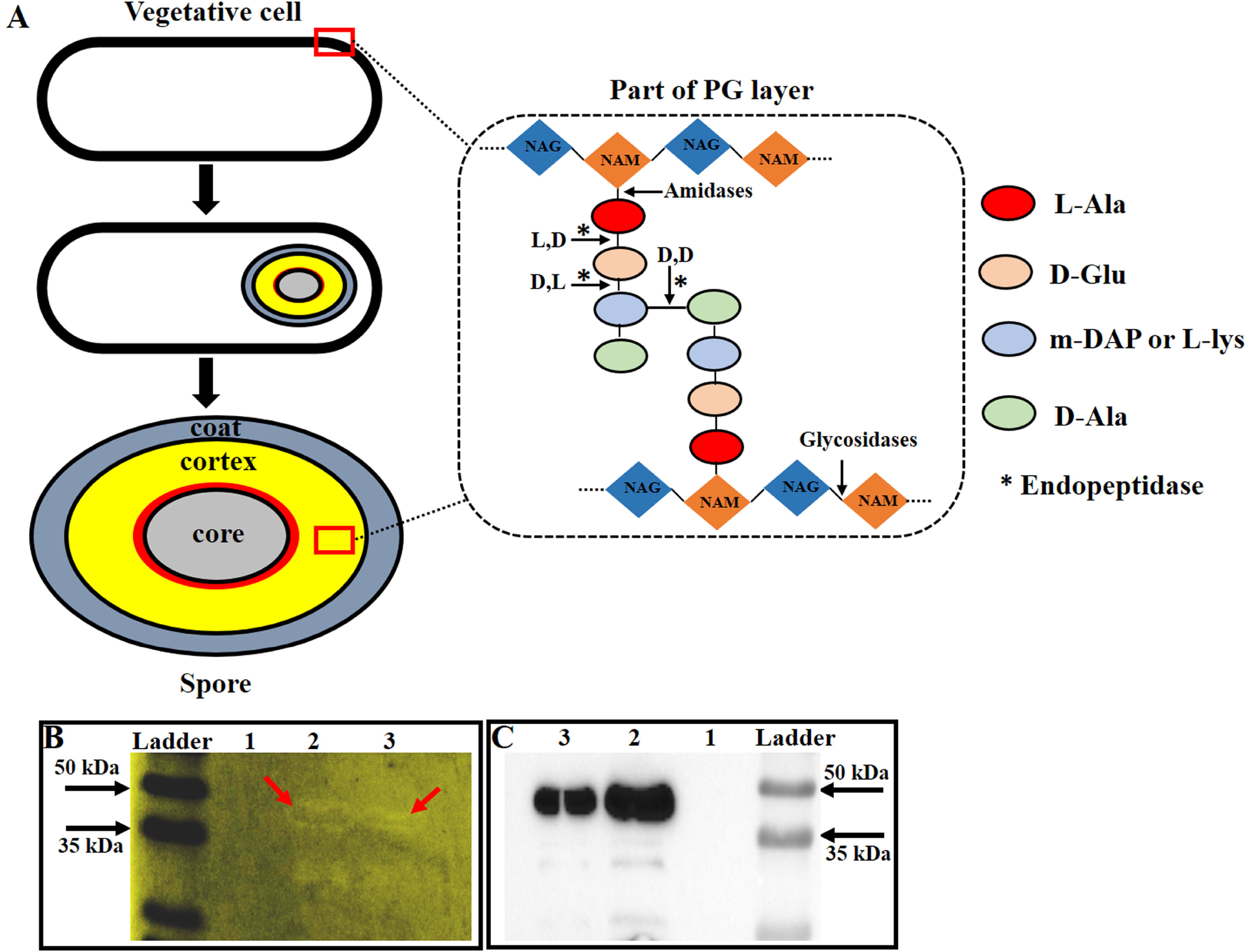
Detection of Cwl0971 hydrolytic activity with zymogram assay. A. Schematic representation of peptidoglycan (PG) layer of *C. difficile* vegetative cells and spores. Glycan strands are formed by β-1,4-linked-N-acetylmuramic acid (NAM) and N-acetylglucosamine (NAG), cross-linked by short peptides attached to NAM. Endopeptidases (*), glycosidases, and amidases cleavage sites were indicated in the black arrow. B. Zymogram assay. Purified Cwl0971 was used to detect hydrolase activity to the *B. subtilis* cell wall. Lane 1 was blank control. The quantity of Cwl0971 loaded in lane 2 was two folds of lane 3. Lysis was visualized as translucent zones appearing in the gel. C. Western blotting. Zymogram was run in parallel with Western blotting detection gels to confirm the proteins migrated at the correct size.

### Effect of Cwl0971 defect on bacterial cell viability

To determine if Cwl0971 could affect *C. difficile* cell autolysis, the Triton X-100 autolysis assay was conducted. No significant difference in bacterial autolysis of R20291 VS R20291-pMTL84153 and R20291Δ0971 VS R20291Δ0971-pMTL84153 was detected (data not shown). As shown in Fig. 3A, R20291Δ0971 lysed at a slower rate than R20291 (*\*P* < 0.05) indicating that the Cwl0971 defect increased bacterial resistance to Triton X-100 and impaired autolysis of R20291. The lysed *C. difficile* cells with Triton X-100 induction were visualized by phase-contrast microscopy and DAPI / PI co-staining (Fig. S1).

**Fig. 3.**
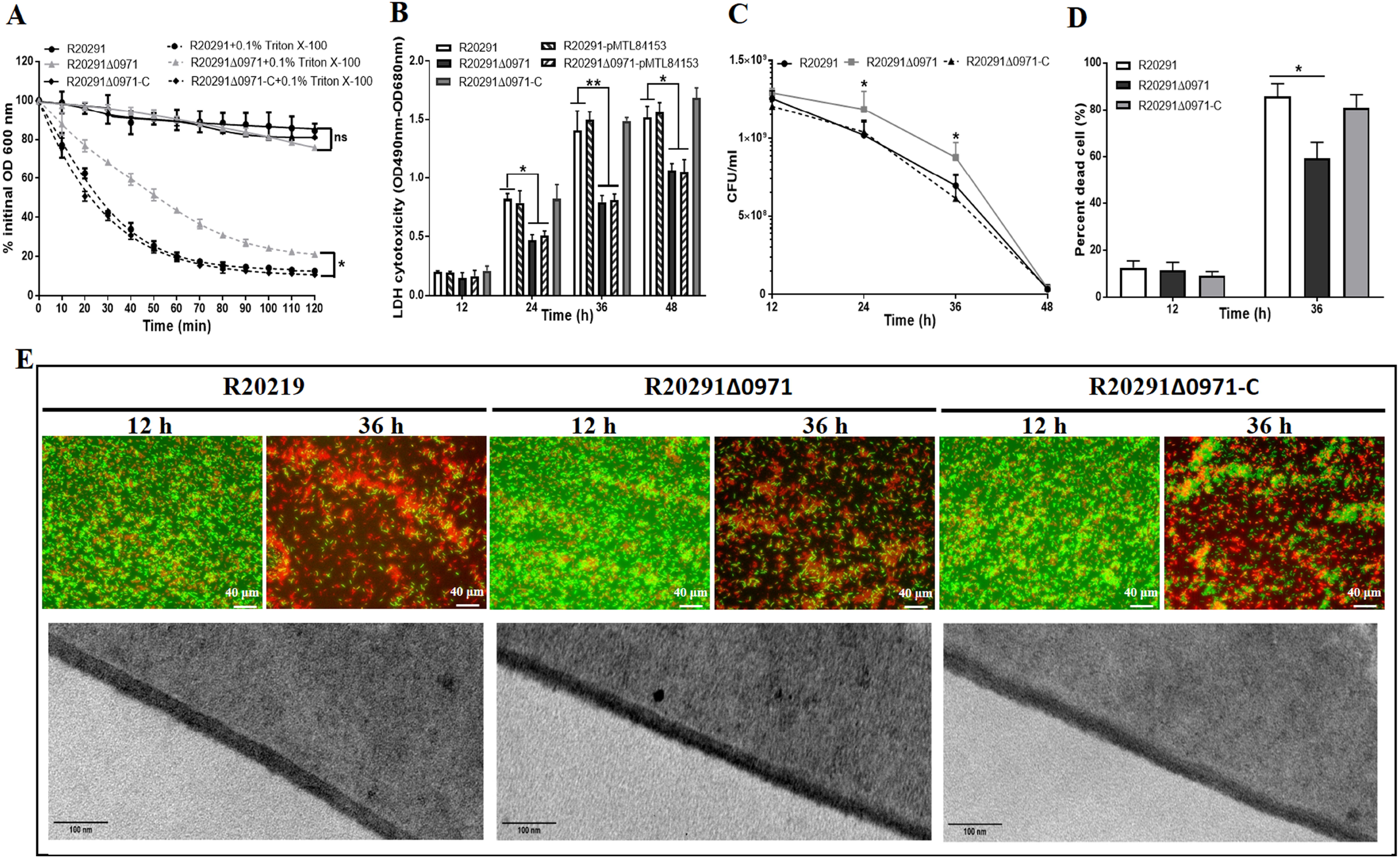
Effect of Cwl0971 defect on cell viability. A. Triton X-100 autolysis assay. The *C. difficile* pellets collected from the log phase were resuspended in 50 mM potassium phosphate buffer with or without 0.1% Triton X-100, and the OD_600_ was detected once every 10 min for 120 min. The lysis percent was shown as % initial OD_600_. B. LDH cytotoxicity assay. LDH concentration of *C. difficile* supernatants at 12, 24, 36, and 48 h post-inoculation was detected with the Pierce™ LDH Cytotoxicity Assay Kit. C. Enumeration of live cells in *C. difficile* cultures at 12, 24, 36, and 48 h post-inoculation on BHIS plates. D. Percent of dead cells in live-dead cell staining slides as shown in (E). E. Cell viability detection by live-dead cell staining and cell wall structure visulization by TEM. *C. difficile* cells were co-dyed with CFDA (green) / PI (red) and monitored by a fluorescent microscope at 40 × magnification. TEM figures were obtained with a JEOL JEM1400 TEM using an accelerating voltage of 80 kV at 120 k × magnification. Experiments were independently repeated thrice. Bars stand for mean ± SEM. One-way analysis of variance (ANOVA) with post-hoc Tukey test was used for statistical significance, statistically significant outcomes are indicted (*\*P* < 0.05, \*\**P* < 0.01).

Lactate dehydrogenase (LDH) is a cytosolic enzyme present in many different cell types. Damage of the plasma membrane can result in a release of LDH into the surrounding cell culture medium. The LDH cytotoxicity of *C. difficile* strains was detected to indicate the potential cell wall permeability and / or cell lysis change. As shown in Fig. 3B, no significant difference of the LDH release can be detected at 12 h post-incubation, while the LDH concentration of R20291Δ0971 was lower than R20291 with a significant difference after 24 h post-incubation (*\*P* < 0.05). Besides, as shown Fig. 1D, the growth curve of parent and mutant strains were similar before 14 h which indicated the similar cell density / number in cultures. These results indicate that the lower LDH release of the mutant is caused by less cell lysis and lower accumulated LDH in the supernatants, rather than caused by the lower cell permeability. To further analyze the effect of Cwl0971 defect on *C. difficile* cell, we analyzed the cell viability through CFU enumeration of *C. difficile* cultures (Fig. 3C) and live-dead cell staining (Fig. 3D and 3E) and performed TEM microscopy to detect cell wall structure (Fig. 3E low panel and Fig. S2). As shown in Fig. 3C, the CFU of the mutant cultures was significantly higher than that of the wild type strain at 24 and 36 h post-incubation (*\*P* < 0.05). For calculation of dead cell percentage in the live-dead staining slides, four areas of cells (>400 cells) on a slide were counted with microscope software (dead bacteria were dyed as the red color with PI, and live bacteria were dyed as the green color with CFDA), and the percent of dead cells accounted in total cells were calculated (Fig. 3D). As shown in Fig. 3D and 3E, a significant reduction of dead cells of R20291Δ0971 was found at 36 h post-incubation (*\*P* < 0.05). In the TEM microscopy, no significant difference in cell wall thickness could be detected among R20291 (28.81 ± 0.98 nm), R20291Δ0971 (31.07 ± 1.50 nm), and R20291Δ0971-C (29.38 ± 0.57 nm). Though we tried many times, we didn’t observe visible changes of *C. difficile* cell wall structure in the mutant by TEM microscopy. Taken together, our data demonstrated that Cwl0971 impairs the bacterial autolysis which can increase the cell viability of R20291Δ0971.

### Effects of Cwl0971 defect on toxin release

To evaluate the effect of 0971 gene deletion on toxin production, the transcription of toxin genes (*tcdA* and *tcdB*) and the toxin concentration of culture supernatants at 12, 24, 36, and 48 h post-inoculation were firstly analyzed by RT-qPCR and ELISA. As shown in Fig. 4A and 4D, no significant difference in toxin transcription of R20291 VS R20291Δ0971 was detected, while Fig. 4B and 4E showed that the concentration of TcdA and TcdB of R20291Δ0971 was significantly lower than that of R20291 after 36 h post-inoculation (\**P* □ 0.05). The TcdA concentration of R20291Δ0971 decreased by 26.5% (\**P* □ 0.05) at 36 h and 25.0% (\**P* □ 0.05) at 48 h (Fig. 4B), and the TcdB concentration of R20291Δ0971 decreased by 15.1% (\**P* □ 0.05) at 36 h and 11.8% (\**P* □ 0.05) at 48 h (Fig. 4E). To evaluate the total toxin production in *C. difficile* cells, the intracellular toxin concentration was determined by ELISA as well (Fig. 4C and 4E). Fig. 4C and 4E showed that no significant difference in total toxin production of R20291 VS R20291Δ0971 was detected, which was consistent with the toxin gene transcription data. Taken together, our data suggested that Cwl0971 does not affect toxin transcription nor total toxin production, but rather facilitates toxin release by affecting bacterial autolysis.

**Fig. 4.**
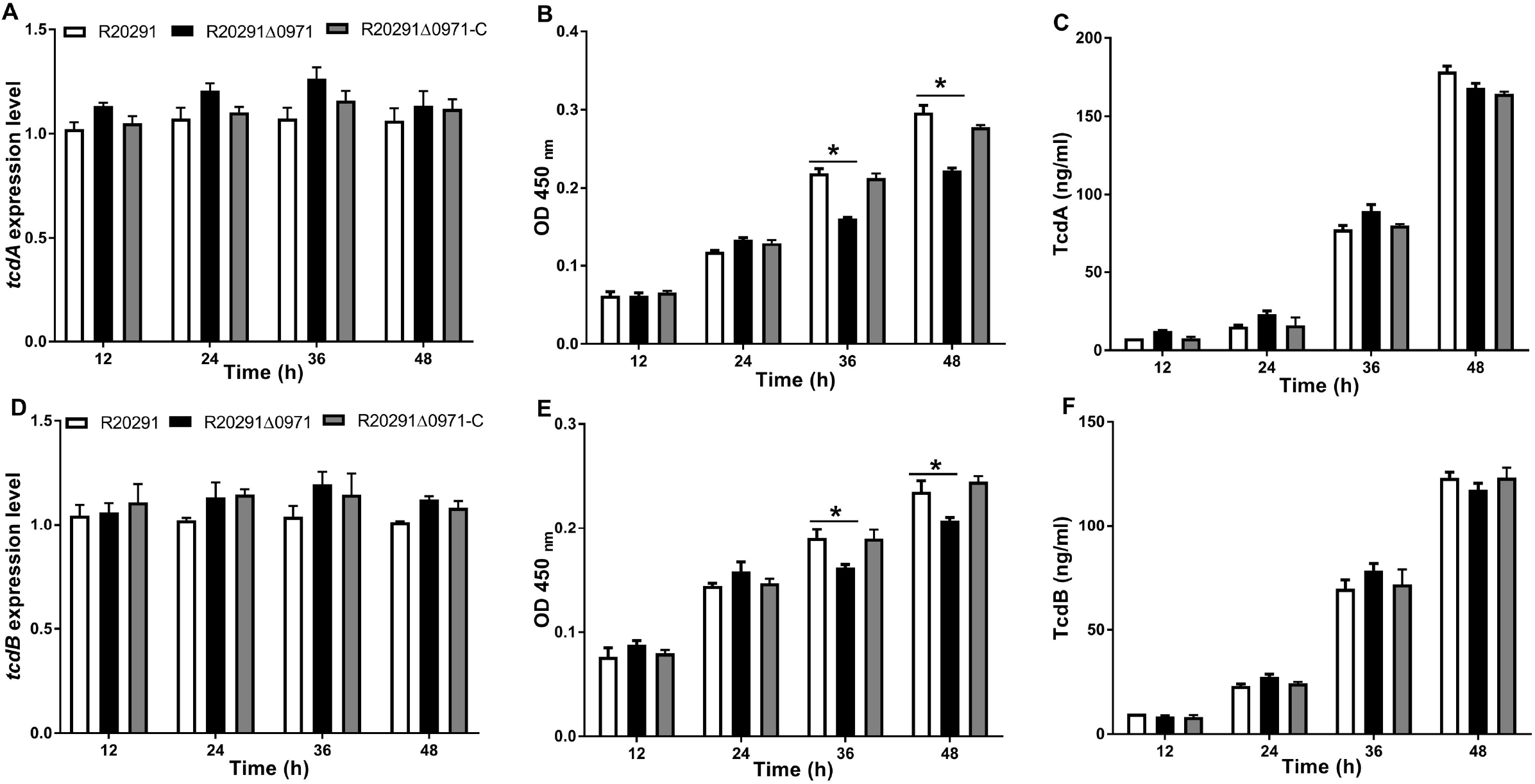
Effect of Cwl0971 defect on toxin release. *C. difficile* cultures were collected at 12, 24, 36, and 48 h post-incubation for toxin production analysis. A. *tcdA* expression on transcription level. B. TcdA concentration in *C. difficile* culture supernatants detected by ELISA. C. Intracellular TcdA concentration detected by ELISA. D. *tcdB* expression on transcription level. E. TcdB concentration in *C. difficile* culture supernatants. F. Intracellular TcdB concentration detected by ELISA. Experiments were independently repeated thrice. Bars stand for mean ± SEM. One-way analysis of variance (ANOVA) with post-hoc Tukey test was used for statistical significance to compare toxin production in different *C. difficile* strains, statistically significant outcomes are indicted (*\*P* < 0.05).

### Effects of Cwl0971 defect on sporulation, germination, and cell length

*C. difficile* strains were cultured in Clospore media for 120 h for sporulation ratio analysis, followed by enumerating the CFU of *C. difficile* cultures on BHIS plates with or without 0.1% TA. As shown in Fig. 5A, the sporulation ratio of R20291Δ0971 decreased by 63.0% (\*\**P* □ 0.01) compared to that of R20291 at 120 h post-incubation. To further visualize the spores and determine the sporulation ratio, phase-contrast microscopy was performed. As shown in Fig. 5B and Fig. S3, R20291Δ0971 showed a significant reduction in sporulation compared to that of wild type strain (\**P* □ 0.05). Expression of sporulation regulation sigma genes *spo0A*, *sigE*, *sigF*, and *sigG* was detected by RT-qPCR (Fimlaid and Shen, 2015) (Fig. 5C). Results showed that the transcription of *spo0A*, *sigE*, *sigF*, and *sigG* in R20291Δ0971 decreased by 38.2% (\**P* □ 0.05), 71.0% (\*\*\*\**P* □ 0.0001), 35.3% (\**P* □ 0.05), and 42.3% (\**P* □ 0.05) compared to that of R20291, respectively.

**Fig. 5.**
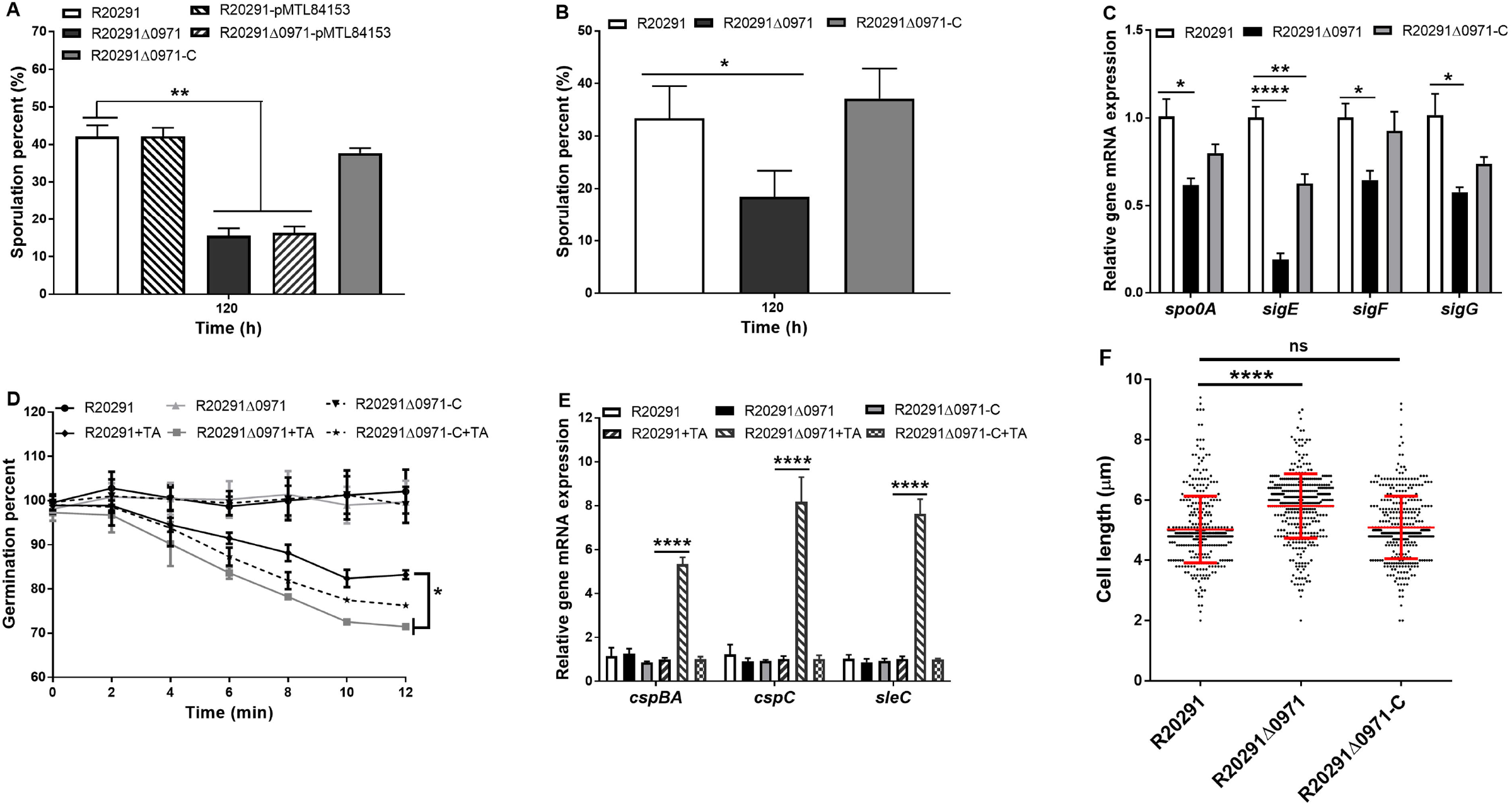
Effect of Cwl0971 defect on sporulation, germination, and cell length. A. Sporulation assay by plating. *C. difficile* strains were cultured in Clospore media for 120 h, following the cultures were 10-fold diluted and plated on BHIS plates with 0.1% TA to detect sporulation ratio. B. Sporulation assay by phase-contrast microscopy. *C. difficile* strains were cultured in Clospore media for 120 h. Then, the spores were numerated by phase-contrast microscopy. At least three fields of view for each strain were acquired to calculate the percentage of spores. C. Essential sporulation related genes expression during *C. difficile* sporulation process. Transcription of *spo0A*, *sigE*, *sigF*, and *sigG* were detected by RT-qPCR at 36 h post-inoculation. D. Germination assay. The purified spores were diluted to OD_600_ of 1.0 in the germination buffer to detect the germination ratio. The boiled spores which were heated at 100 □ for 20 min were used as a negative control (without TA) for germination analysis. E. Germination essential genes expression during the spores germination process. Transcription of *cspBA*, *cspC*, and *sleC* were detected by RT-qPCR in the germination buffer with TA or without TA added. F. *C. difficile* vegetive cell length was measured by phase-contrast microscopy at 6 h and 10 h of post-inoculation. Experiments were independently repeated thrice. Bars stand for mean ± SEM. One-way analysis of variance (ANOVA) with post-hoc Tukey test was used for statistical significance, statistically significant outcomes are indicted (\**P* < 0.05, \*\**P* < 0.01, \*\*\**P* < 0.001, \*\*\*\**P* < 0.0001).

The germination ratio of *C. difficile* spores was evaluated as well. Fig. 5D showed that the 0971 gene deletion significantly increased R20291 spores germination (\**P* □ 0.05). The increase of germination ratio in R20291Δ0971 prompted us to identify if the 0971 gene deletion could affect the expression of essential germination regulation gene *cspBA*, *cspC*, and *sleC* (Paredes-Sabja et al., 2008; Francis et al., 2013; Francis et al., 2015; Bhattacharjee et al., 2016). Transcription of the germination regulation genes was measured by RT-qPCR (Fig. 5E). Interestingly, the transcription of *cspBA*, *cspC*, and *sleC* in R20291Δ0971 increased 4.3 (\*\*\*\**P* □ 0.0001), 7.2 (\*\*\*\**P* □ 0.0001), and 6.6 (\*\*\*\**P* □ 0.0001) folds compared to that of R20291, respectively, indicating a putative effect of 0971 gene defect on *cspBA*, *cspC*, and *sleC* transcription, while the detailed correlation remains to be uncovered.

Endopeptidase can remodel the bacterial cell wall through hydrolyzing the cell wall peptidoglycan to affect cell separation and cell length (Vollmer et al., 2008). The cell length and separation of R20291Δ0971 were determined by phase-contrast microscopy (Fig. 5F, Fig. S2, and Fig. S4). Results showed that the cell length of R20291Δ0971 (5.805 ± 1.068 μm, \*\*\*\**P* □ 0.0001, 399 cells counted) significantly increased compared to that of R20291 (5.023 ± 1.105 μm, 408 cells counted), and the cell length can be recovered when the 0971 gene was complemented in R20291Δ0971 (5.094 ± 1.036 μm, 402 cells counted) (Fig. 5G). While the cell length was affected by 0971 gene deletion, we didn’t detect the significant difference in cell separation and septa of R20291 VS R20291Δ0971 (Fig. S4 and Fig. S2). Taken together, our data showed that Cwl0971 defect can affect *C. difficile* sporulation and germination and increase *C. difficile* cell length.

### Evaluation of Cwl0971 defect on bacterial virulence in the CDI mouse model

To evaluate the effect of 0971 gene deletion on *C. difficile* virulence *in vivo*, the mouse model of CDI was used. Twenty mice (n=10 per group) were orally challenged with R20291 or R20291Δ0971 spores (1 × 10^6^ spores / mouse) after antibiotic treatment. As shown in Fig. 6A, the R20291Δ0971 infection group lost less weight at days post challenge 3 (\**P* □ 0.05) compared to the R20291 infection group. Fig. 6B showed that only 20% of mice succumbed to severe disease within 4 days in the R20291Δ0971 infection group compared to 50% mortality in the R20291 infection group (no significant difference with log-rank analysis). Meanwhile, 100% of mice developed diarrhea in the R20291 infection group versus 80% in the R20291Δ0971 infection group at days post challenge 2 (Fig. 6C). As shown in Fig. 6D, the CFU of the R20291Δ0971 infection group decreased in the fecal shedding samples at days post challenge 1, 2, and 4 (\**P* □ 0.05) compared to the R20291 infection group.

**Fig. 6.**
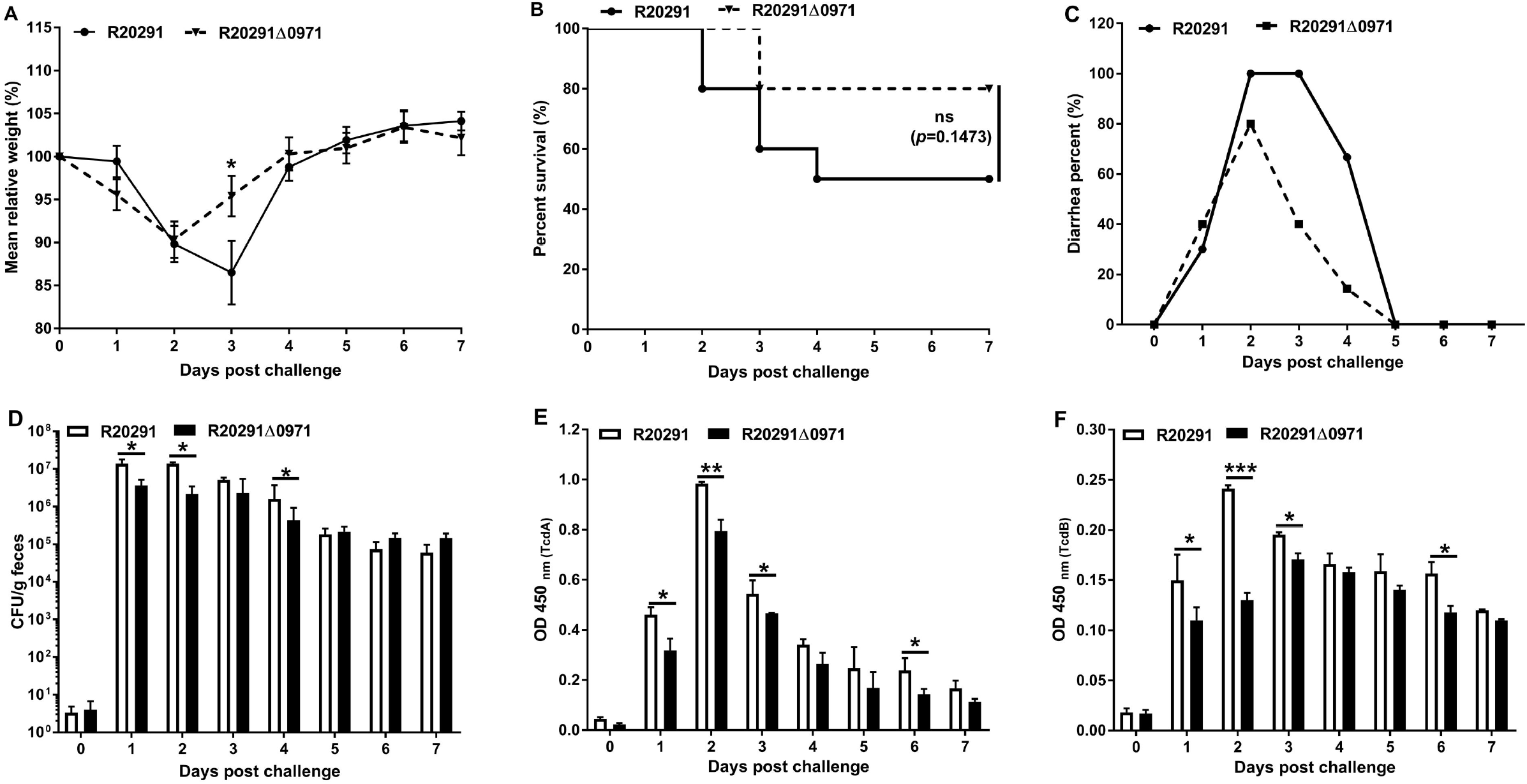
Evaluation of Cwl0971 defect on *C. difficile* virulence in mice. A. Mean relative weight changes. B. Survival curves. C. Diarrhea percentages. D. *C. difficile* spore numbers in feces. E. TcdA level of fecal sample. F. TcdB level of fecal sample. Bars stand for mean ± SEM. Student’s unpaired *t*-test was used for statistical significance, statistically significant outcomes are indicted (\**P* < 0.05, \*\**P* < 0.01, \*\*\**P* < 0.001). Animal survivals were analyzed by Kaplan-Meier survival analysis with a log-rank test of significance.

To evaluate the toxin level in the gut, the titer of TcdA and TcdB in the feces was measured. In comparison with the R20291 infection group, the TcdA and TcdB of the R20291Δ0971 infection group decreased significantly at days post challenge 1 (\**P* □ 0.05), 2 (TcdA \*\**P* □ 0.01, TcdB \*\*\**P* □ 0.001), 3 (\**P* □ 0.05), and 6 (\**P* □ 0.05) (Fig. 6E and 6F). Taken together, our results indicated that the Cwl0971 defect impairs R20291 pathogenicity.

## Discussion

In this study, we identified and characterized a novel peptidoglycan hydrolase (putative endopeptidase) Cwl0971 from *C. difficile* R20291. Our data suggest that Cwl0971 has a hydrolytic activity for bacterial cell wall and is involved in different cellular processes of *C. difficile* such as cell viability, sporulation, and spores germination. Notably, the defect of Cwl0971 decreased bacteriolysis, toxin release, and sporulation, and decreased fitness over the wild strain in the mouse infection model, indicating that Cwl0971 could be a new potential target for the development of antimicrobials against *C. difficile.*

C40 family peptidase/NlpC/P60 protein usually contains a conserved Cys-His-His catalytic triad that appears to be specific to murein tetrapeptide, and typically cleaves the linkage between D-Glu and L-diaminopimelic acid (L-DPA) (or Lys) within peptidoglycan stem peptides that was involved in cell wall hydrolysis during cell growth or cell lysis (Anantharaman and Aravind, 2003). Cwl0971 is composed of 3 SH3_3 cell wall / substrate-binding domains in the N-terminus and an NlpC/P60 (pfam00877) domain in the C-terminus with endopeptidase activity predicated through bioinformatics analysis with NCBI’s Conserved Domain Database (CDD). SH3 (Src Homology 3) domains are protein interaction domains that bind proline-rich ligands with moderate affinity and selectivity, preferentially to PxxP motifs (Xu et al., 2015). SH3 domains containing proteins play versatile and diverse roles in the cell, including regulation of enzymes, changing the subcellular localization of signaling pathway components, and mediating the formation of multiprotein complex assemblies (https://www.ncbi.nlm.nih.gov/Structure/cdd/cddsrv.cgi?ascbin=8&maxaln=10&seltype=2&uid=388381). Though we didn’t identify the endopeptidase activity and detail cleavage sites of Cwl0971, the hydrolytic activity of Cwl0971 was confirmed by zymography in our study (Fig. 2B). Notably, the homolog protein CD1135 (CwlA) in *C.difficile* 630 was recently characterized as a γ-D-Glu-mDAP-endopeptidase by using RP-HPLC (Garcia-Garcia et al., 2020).

Peptidoglycan (PG), a dynamic macromolecule, is a primary cell wall constituent of Gram-positive bacteria and is constantly remodeled to enable bacterial growth and differentiation. Peptidoglycan hydrolases (PGHs) contribute to PG plasticity and play a key role in maintaining the cell wall shape through hydrolyzation of PG bonds. Among them, endopeptidase is one major peptidoglycan hydrolase involved in several cellular processes (Layec et al., 2008; Vollmer et al., 2008). Endopeptidase can be subclassified as D,D-endopeptidase, D,L-endopeptidase, and L,D-endopeptidase depending on their hydrolytic specificity. Among them the D,L-endopeptidases usually cleave the bond between D-Glu (position 2) and DPA (position 3) of the peptide and are classified as two different families based on Zn^2+^ dependence (Hourdou et al., 1992). CwlS, CwlO, LytE, and LytF from *B. sphaericus* and p60 from *L. monocytogenes* which belong to the family II D,L-endopeptidases have been shown to regulate cell autolysis and division (Margot et al., 1998; Ohnishi et al., 1999; Pilgrim et al., 2003; Yamaguchi et al., 2004; Fukushima et al., 2006). YqgT, the Zn^2+^ dependent family I D,L-endopeptidases from *B. sphaericus*, was studied as well (Hourdou et al., 1993). Other endopeptidases, CwlK and LytH, were also reported to be involved in bacterial physiology (Horsburgh et al., 2003; Fukushima et al., 2007). Recently, one study published on bioRxiv by Garcia-Garcia et. al (Garcia-Garcia et al., 2020) showed that CwlA (the homologue of Cwl0971 in CD630) could modulate *C.difficile* cell division through serine/threonine kinase (PrkC)-dependent phosphorylation, suggesting a novel regulation mechanism for cell wall hydrolysis in *C. difficile* 630. Though the cell length of R20291Δ0971 was significantly longer than that of R20291 (Fig. 5F, Fig. S2, and Fig. S4), we didn’t observe the significant difference in cell separation between mutant and wild type strain. Besides, the transcription of 0971 gene peaked at log-phase followed by a reduction at stationary-phase, and increased again at decline-phase, indicating the potential critical roles of Cwl0971 at multiple stages in R20291growth (Fig. 1E). These results prompted us to examine the other effects of Cwl0971 defect on R20291 physiology, especially toxin release, sporulation, and germination.

Two exotoxins (TcdA and TcdB) have been identified as the major virulence factors of *C. difficile* (Kuehne et al., 2010). Several genes, such as *tcdC*, *tcdE*, and *tcdR*, are involved in the regulation of toxin gene expression (Mani et al., 2002; Dupuy et al., 2008; Govind and Dupuy, 2012; Govind et al., 2015). TcdA and TcdB are the large clostridial glucosylating family toxins that are secreted without signal peptide (Popoff and Bouvet, 2009). Except for reported regulation genes, toxin release is also associated with *C. difficile* cell autolysis. Recently, Cwp19 has been identified as a novel lytic transglycosylase that can generate muropeptide (Wydau-Dematteis et al., 2018). Cwp19 was reported to facilitate toxin release through bacteriolysis, which indicated that TcdE and bacteriolysis were the coexisting mechanisms for toxin release. Cwp22, a novel peptidoglycan cross-linking enzyme, was reported to be involved in *C. difficile* cell wall integrity and toxin release as well (Zhu et al., 2019). In this study, our data clearly show that Cwl0971 affects cell viability by impairing cell autolysis in *C. difficile* (Fig. 3). We further assessed toxin production of the mutant and parent strains (Fig. 4). Lower toxin concentration in the supernatant of the mutant after 36 h post-inoculation was detected, while toxin gene transcription and intracellular total toxin concentration were at a similar level between R20291 and R20291Δ0971. Our data suggest that Cwl0971 contributes to toxin release by affecting bacteriolysis as well.

PGHs have been reported to be involved in several cellular processes (Layec et al., 2008; Vollmer et al., 2008). Haiser et al. (Haiser et al., 2009) demonstrated that cell wall hydrolase RpfA, SwlA, SwlB, and SwlC play critical roles at multiple stages in *Streptomyces coelicolor* growth and development. The deletion of each of these four hydrolase genes could impair bacteria heat resistance, vegetative growth, spore formation, and spore germination. To cause disease, *C. difficile* spores must germinate into the vegetative cells. In *C. difficile*, sporulation and germination play key roles in *C. difficile* pathogenesis, transmission, and persistence of CDI (Haiser et al., 2009; Gil et al., 2017; Zhu et al., 2018). So far, several PGHs in *C. difficile* 630 have been reported, which could be involved in bacteriolysis, engulfment, sporulation, heat-resistant spore formation, and germination (Dhalluin et al., 2005; Layec et al., 2008; Gutelius et al., 2014; Xu et al., 2014; Fimlaid et al., 2015). These reported studies prompted us to examine the effects of 0971 gene deletion on some other *C. difficile* phenotypes, especially sporulation and germination. We found that R20291Δ0971 exhibited a significant defect in sporulation (Fig. 5A and 5B). The master sporulation regulator Spo0A plays a critical role in *C. difficile* sporulation by regulating sporulation-specific RNA polymerase sigma factors, especially σ^E^, σ^F^, σ^G^, and σ^K^ (Fimlaid and Shen, 2015). Therefore, we analyzed the transcription of *spo0A*, *sigE*, *sigF*, and *sigG* by RT-qPCR. The significant decrease of sigma factors expression may partially explain the sporulation decrease of R20291Δ0971 (Fig. 5C), while how Cwl0971 affects sigma factors expression remains to be further studied. Spores germination of R20291Δ0971 was also compared to that of R20291. We found that R20291Δ0971 displayed a significantly increased germination ratio in comparison with R20291 (Fig. 5D). *C*. *difficile* packages three essential subtilisin-like serine proteases proteins, CspA, CspB, and CspC into spores for spores germination (Paredes-Sabja et al., 2008; Francis et al., 2013; Francis et al., 2015; Bhattacharjee et al., 2016). Meanwhile, the cortex hydrolase SleC also plays a critical role in cortex degradation for spores germination (Francis and Sorg, 2016). As expected, the transcription of *cspBA*, *cspC*, and *sleC* in R20291Δ0971 significantly increased compared to that of R20291 in the germination process (Fig. 5E). Our data suggest that Cwl0971 plays a positive effect on sporulation and a negative effect on spores germination, more details need to be uncovered to characterize how Cwl0971 affects these physiological processes.

We also assayed biofilm formation, adhesion, spore heat-resistance, spore structure, and motility of the mutant and wild type strains. We found that Cwl0971 defect impairs biofilm production and spore resistance to heat, while increases cell motility and adhesion (Fig. S5 and Fig. S6). Biofilms contribute to survival, persistence, antimicrobial resistance, colonization, and disease for many pathogens and it has been reported that up to 80% of bacterial infections were linked to biofilms (Flemming and Wingender, 2010). Previous studies have highlighted that the flagella (motility) of *C. difficile* play an important role in biofilm formation and bacteria adherence to host (Tasteyre et al., 2001), but the results from different groups are controversial. In *C. difficile* 630Δerm, previous reports have shown that the *fliC* mutation (motility defect) could enhance toxin production and bacterial adherence to the host (Dingle et al., 2011; Aubry et al., 2012; Baban et al., 2013a). While in R20291, the *fliC* mutation decreased biofilm formation and adherence to Caco-2 cells (Baban et al., 2013a; Ethapa et al., 2013). We found that the biofilm production of R20291Δ0971 significantly decreased, while the adhesion and mobile ability of R20291Δ0971 significanlty increased (Fig. S5 and S6). The complex relationship between biofilm formation, motility, and adhesin requires further study to fully understand their coordinated mechanism. Collectively, our data suggest that Cwl0971 defect affects different cellular processes involved in *C. difficile* pathogenesis (Fig. 6).

## Experimental procedures

### Comparative genomic analysis of *C. difficile* genomes

Using the Vaxign reverse vaccinology tool (He et al., 2010), we systematically analyzed all proteins in the genome of *C. difficile* R20291 in terms of cellular localization, adhesin probability, transmembrane helices, sequence conversation with the genomes of other 12 *C. difficile* strains, sequence similarity to human and mouse proteins and protein length. These other 12 strains are strains 630, BI1, ATCC 43255, CD196, CIP 107932, QCD-23m63, QCD-32g58, QCD-37×79, QCD-63q42, QCD-66c26, QCD-76w55, and QCD-97b34. Protein-conserved domain analysis was performed using the NCBI’s CDD.

### Bacteria, plasmids, and culture conditions

Table 1 lists the strains and plasmids used in this study. *C. difficile* strains were cultured in BHIS (brain heart infusion broth supplemented with 0.5% yeast extract and 0.1% L-cysteine, and 1.5% agar for agar plates) at 37 □ in an anaerobic chamber (90% N_2_, 5% H_2_, 5% CO_2_). For spore preparation, *C. difficile* strains were cultured in Clospore media and purified as described (Perez et al., 2011). *Escherichia coli* DH5α, *E. coli* HB101/pRK24, and *E. coli* BL21 (DE3) were grown aerobically at 37 □ in LB media (1% tryptone, 0.5% yeast extract, 1% NaCl). Among them, *E. coli* DH5α was used as a cloning host, *E. coli* HB101/pRK24 was used as a conjugation donor host, and *E. coli* BL21 (DE3) was used as a protein expression host. Antibiotics were added when needed: for *E. coli*, 15 μg ml^−1^ chloramphenicol, 50 μg ml^−1^ kanamycin; for *C. difficile*, 15 μg ml^−1^ thiamphenicol, 250 μg ml^−1^ D-cycloserine, 50 μg ml^−1^ kanamycin, 8 μg ml^−1^ cefoxitin, and 500 ng ml^−1^ anhydrotetracycline.

**Table 1.**
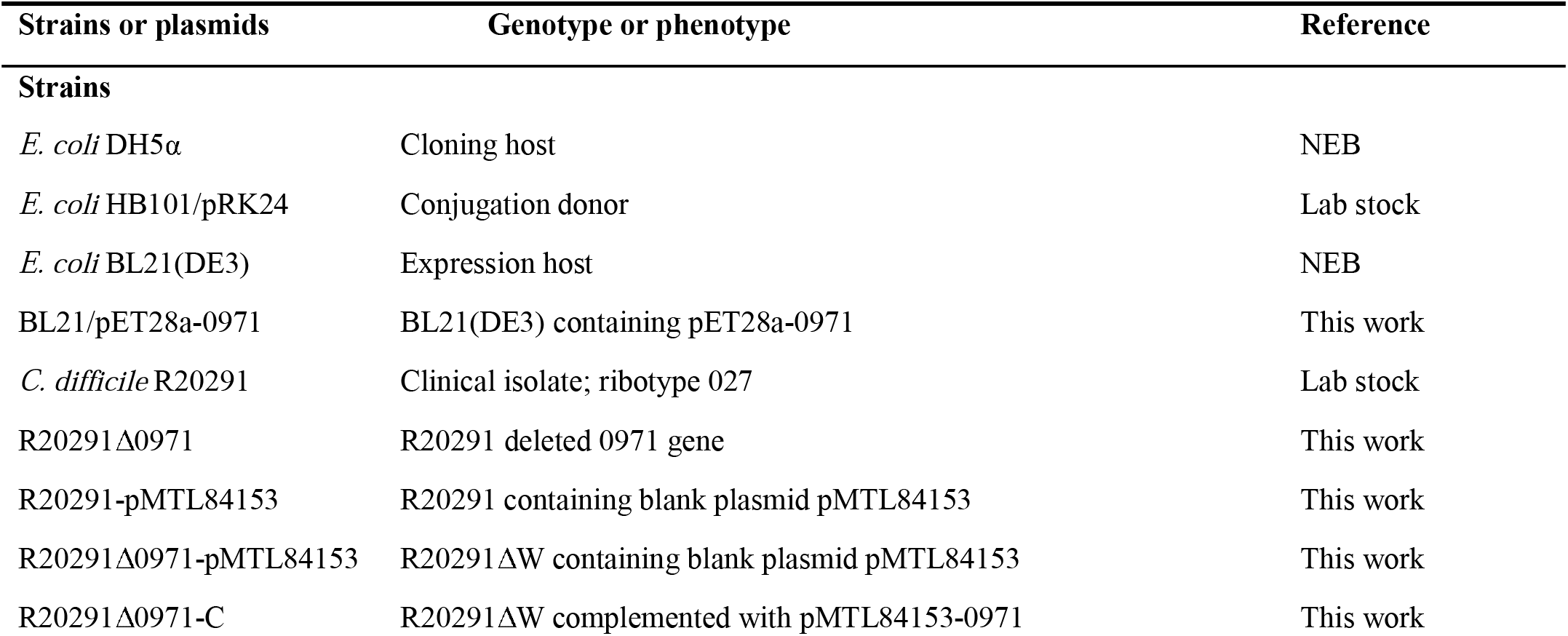
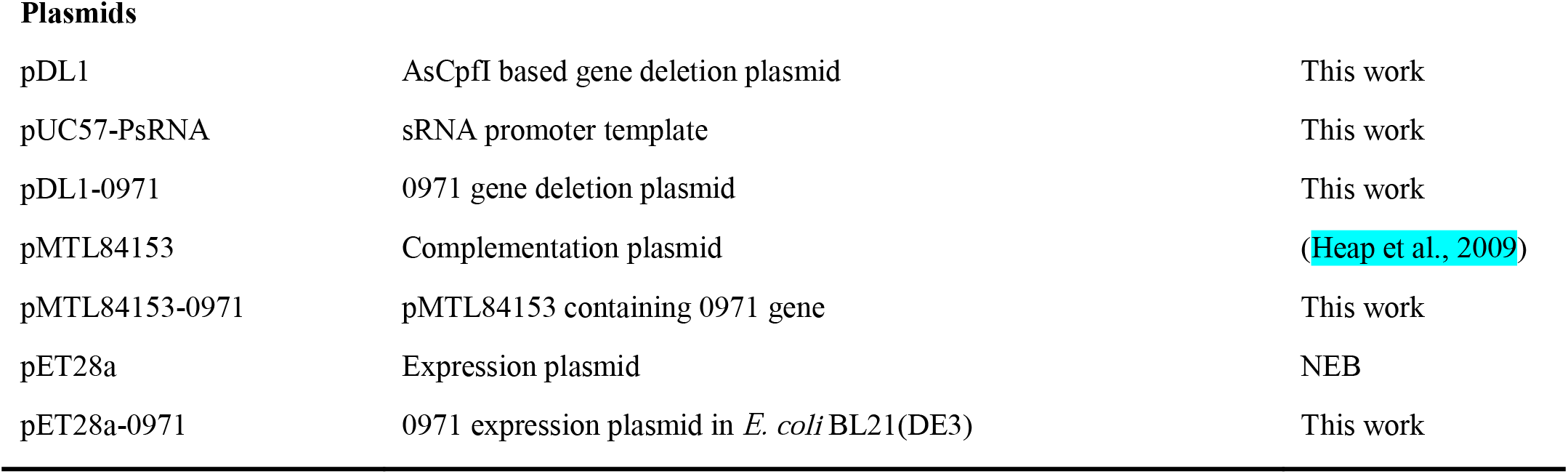
Bacteria and plasmids utilized in this study.

| Strains or plasmids | Genotype or phenotype | Reference |
| --- | --- | --- |
| **Strains** |  |  |
| *E. coli* DH5α | Cloning host | NEB |
| *E. coli* HB101/pRK24 | Conjugation donor | Lab stock |
| *E. coli* BL21(DE3) | Expression host | NEB |
| BL21/pET28a-0971 | BL21(DE3) containing pET28a-0971 | This work |
| *C. difficile* R20291 | Clinical isolate; ribotype 027 | Lab stock |
| R20291Δ0971 | R20291 deleted 0971 gene | This work |
| R20291-pMTL84153 | R20291 containing blank plasmid pMTL84153 | This work |
| R20291Δ0971-pMTL84153 | R20291ΔW containing blank plasmid pMTL84153 | This work |
| R20291Δ0971-C | R20291ΔW complemented with pMTL84153-0971 | This work |
| Plasmids |  |  |
| pDL1 | AsCpfI based gene deletion plasmid | This work |
| pUC57-PsRNA | sRNA promoter template | This work |
| pDL1-0971 | 0971 gene deletion plasmid | This work |
| pMTL84153 | Complementation plasmid | (Heap et al., 2009) |
| pMTL84153-0971 | pMTL84153 containing 0971 gene | This work |
| pET28a | Expression plasmid | NEB |
| pET28a-0971 | 0971 expression plasmid in *E. coli* BL21(DE3) | This work |

### DNA manipulations and chemicals

DNA manipulations were carried out according to standard techniques (Chong, 2001). Plasmids were conjugated into *C. difficile* as described earlier (Heap et al., 2010). The DNA markers, protein markers, PCR product purification kit, DNA gel extraction kit, restriction enzymes, cDNA synthesis kit, and SYBR Green RT-qPCR kit were purchased from Thermo Fisher Scientific (Waltham, USA). PCRs were performed with the high-fidelity DNA polymerase NEB Q5 Master Mix, and PCR products were assembled into target plasmids with NEBuilder HIFI DNA Assembly Master Mix (New England, UK). Primers (Supporting Information Table S1) were purchased from IDT (Coralville, USA). All chemicals were purchased from Sigma (St. louis, USA) unless those stated otherwise.

### Construction of 0971 deletion mutant and complementation strains

The Cas12a (AsCpfI) based gene deletion plasmid pDL-1 was constructed and used for *C. difficile* gene deletion (Hong et al., 2018). Target sgRNAs were designed with an available website tool (http://big.hanyang.ac.kr/cindel/), and the off-target prediction was analyzed on the Cas-OFFinder website (http://www.rgenome.net/cas-offinder/). The sgRNA and homologous arms (up and down) were assembled into pDL-1. Two sgRNAs targeting the 0971 gene were designed and used for gene deletion plasmid construction in *C. difficile*. Briefly, the gene deletion plasmid was constructed in the cloning host *E. coli* DH5α and was transformed into the donor host *E. coli* HB101/pRK24, and subsequently was conjugated into R20291. Potential successful transconjugants were selected with selective antibiotic BHIS-TKC plates (15 μg ml^−1^ thiamphenicol, 50 μg ml^−1^ kanamycin, 8 μg ml^−1^ cefoxitin). The transconjugants were cultured in BHIS-Tm broth (15 μg ml^−1^ thiamphenicol) to log phase, then the subsequent cultures were plated on induction plates (BHIS-Tm-ATc: 15 μg ml^−1^ thiamphenicol and 500 ng ml^−1^ anhydrotetracycline). After 24 - 48 h incubation, 20 - 40 colonies were selected and used as templates for colony PCR test with check primers 1-F/R and 2-F/R. The correct gene deletion colony was subcultured into BHIS broth without antibiotics and was passaged several times for plasmid cure to get a clean gene deletion mutant R20291Δ0971. The R20291Δ0971 genome was isolated and used as the template for PCR test with 1-F/R and 2-F/R primers, and the PCR products were sequenced to confirm the correct gene deletion.

The 0971 gene was amplified with primer 3-F/R and assembled into *Sac*I-*Bam*HI digested pMTL84153 plasmid, yielding the complementation plasmid pMTL84153-0971, and was subsequently conjugated into R20291Δ0971, yielding R20291Δ0971/pMTL84153-0971 (shorted as R20291Δ0971-C). The blank plasmid pMTL84153 was also conjugated into R20291 and R20291Δ0971 as negative controls, respectively.

### Purification and hydrolytic activity assay of Cwl0971

The 0971 gene encoding the predicted binding and catalytic domains (without signal peptide) was PCR amplified with primer 4-F/R and assembled into *Nco*I-*Xho*I digested pET28a expression plasmid. 6×His tag was fused into the N-terminal of Cwl0971 for protein purification with Ni^2+^ affinity chromatography column. The protein was purified as described earlier (Peng et al., 2018).

The hydrolytic activity of purified Cwl0971 was evaluated with a zymogram assay (Leclerc and Asselin, 1989). Briefly, cell wall material (0.1%, wt/vol, 200 μl of a 50 mg ml^−1^ autoclaved *Bacillus subtilis* cell suspension was added) was incorporated into SDS polyacrylamide separating gels. Following constant voltage electrophoresis (100 V) at 4 □, gels were washed 3 times with water for 10 minutes at room temperature, then transferred to a renaturing buffer (20 mM Tris, 50 mM NaCl, 20 mM MgCl_2_, 0.5% Triton X-100, pH 7.4) and washed gently overnight at 37 □. The renatured Cwl0971 appeared as slightly clear bands on an opaque background. The contrast was enhanced by staining the gels with 0.1% (wt/vol) methylene blue, 0.01% (wt/vol) KOH for 2 h. Zymogram was run in parallel with Western blot detection gels to confirm that the proteins migrated at the correct size.

### Growth profile, cell autolysis, LDH cytotoxicity, and cell viability assay

*C. difficile* strains were incubated to an optical density of OD_600_ of 0.8 in BHIS and were diluted to an OD_600_ of 0.2. Then, 1% of the culture was inoculated into fresh BHIS, followed by measuring OD_600_ for 34 h.

To determine Triton X-100 induced autolysis, *C. difficile* strains were cultured to an OD_600_ of 0.8 to log phase, and then 5 ml of each culture was collected and washed with 50 mM potassium phosphate buffer (pH 7.0). The pellets were resuspended in a final volume of 2.5 ml of 50 mM potassium phosphate buffer with or without 0.1% of Triton X-100. Afterward, the bacteria were incubated anaerobically at 37 °C, and the OD_600_ was detected once every 10 min for 120 min. The lysis percent was shown as % initial OD_600_. Meanwhile, the autolysed *C. difficile* cells induced by Triton X-100 at 30 min of post-induction were visualized by DAPI / PI staining.

For LDH cytotoxicity analysis, the supernatants from different strains were collected and filtered with 0.22 μm filters used for LDH concentration detection with the Pierce™ LDH Cytotoxicity Assay Kit (Thermo Fisher, USA) according to the instructions of the manufacturer. For cell viability analysis, the live *C. difficile* cells in culture media were enumerated at 24, 36, and 48 h post-inoculation on BHIS plates. Meanwhile, the live-dead cell staining was performed (Zhu et al., 2019). Briefly, 12 and 36 h post incubated *C. difficile* strains were collected and cell number was normalized to 10^8^ CFU/ml, respectively. Then 1 ml of each strain cultures was centrifuged at 4 □, 5000×*g* for 10 min, and washed 3 times with PBS. Afterward, the bacteria were resuspended in 100 μl of 0.1 mM sodium phosphonate buffer. The chemical 5(6)-CFDA (5- (and −6)-Carboxyfluorescein diacetate) was used to dye live *C. difficile*, and the propidium iodide (PI) was used to dye dead bacteria. The final concentration of 50 mM 5(6)-CFDA and 200 ng ml^−1^ of PI were used to co-dye *C. difficile* strains. *C. difficile* cells were incubated at 4 □ overnight for monitoring under a fluorescence microscope. The CFDA and PI were excited at 495 nm and 538 nm, respectively.

To further compare the vegetative cell wall and spore structure of mutant and parent *C. difficile* strains, TEM analysis was performed. The TEM specimens were prepared according to the previous method used in *C. difficile* (Baban et al., 2013b; Calderon-Romero et al., 2018), and detected with a JEM-1400 (Jeol Ltd., Tokyo, Japan) transmission electron microscope using an accelerating voltage of 80 kV. Briefly, 1 ml of bacterial cultures or 1×10^6^ spores were collected and fixed with 2.5% glutaraldehyde (GA). After fixation, samples were post-fixed with 1% osmium tetroxide (EMS), following the samples were dehydrated with a graded series of ethanol (30, 50, 70, 85, and 100%) and acetone. The dehydrated samples were further washed with 2 parts acetone to 1 part embedding 812 resin, 1 part acetone and 1 part embedding 812 resin, 1 part acetone 2 parts embedding 812 resin, then in pure 812 resin and incubated overnight at 4 □, followed by fresh 812 resin for sectioning. Ultrathin 90 nm sections were further picked up on the grid and were post-stained with uranyl acetate and were examined with a JOEL JEM1400 transmission electron microscope using an accelerating voltage of 80 kV.

### Toxin expression assay

To evaluate the toxin expression in *C. difficile* strains, 10 ml of *C. difficile* cultures were collected at 12, 24, 36, and 48 h post-incubation. The cultures were adjusted to the same density with fresh BHIS. Then the cultures were centrifuged at 4 □, 8000×*g* for 15 min, supernatants filtered with 0.22 μm filters, and used for secretion toxin analysis by ELISA. The pellets were resuspended in 2 ml of PBS and disrupted by Bead Beating, following centrifugated at 4 □, 8000 × *g* for 15 min. The intracellular total protein concentration was measured by using a BCA protein assay (Thermo Scientific, Suwanee, GA), and normalized to the same protein concentration for intracellular toxin analysis by ELISA. Anti-TcdA (PCG4.1, Novus Biologicals, USA) and anti-TcdB (AI, Gene Tex, USA) were used as coating antibodies for ELISA, and HRP-Chicken anti-TcdA and HRP-Chicken anti-TcdB (Gallus Immunotech, USA) were used as detection antibodies.

For toxin gene transcription analysis, 2 ml of 12, 24, 36, and 48 h post inoculated *C. difficile* cultures were centrifuged at 4 □, 12000×*g* for 5 min, respectively. Then, the total RNA of different strains was extracted with TRIzol reagent. The transcription of *tcdA* and *tcdB* was measured by RT-qPCR with Q-*tcdA*-F/R and Q-*tcdB*-F/R primers, respectively. All RT-qPCRs were repeated in triplicate, independently. Data were analyzed by using the comparative CT (2^−ΔΔCT^) method with 16s rRNA as a control.

### Sporulation, germination, and cell length assay

*C. difficile* germination and sporulation analysis were conducted as reported earlier (Zhu et al., 2019). Briefly, for *C. difficile* sporulation analysis, *C. difficile* strains were cultured in Clospore media for 5 days, and the CFU of cultures were counted on BHIS plates with 0.1% TA (T_0_, total number) or without 0.1% TA (T_1_, vegetative cell) to detect sporulation ratio, respectively. The sporulation ratio was calculated as (T_0_-T_1_)/T_0_. To further visualize the spores and determine the sporulation ratio, the phase-contrast microscopy was performed. Briefly, 1 ml of 120 h post-inoculation cultures was centrifuged and resuspended in 10 μl of fresh media. Followed, slides were prepared by placing 2 μl of the concentrated culture onto a thin layer of 0.7% agarose applied directly to the surface of the slide. Phase-contrast microscopy was performed using a ×100 Ph3 oil immersion objective on a Nikon Eclipse Ci-L microscope. At least three fields of view for each strain were acquired with a DS-Fi2 camera and used to calculate the percentage of spores (the number of spores divided by the total number of spores, prespores, and vegetative cells) from three independent experiments.

For *C. difficile* germination assay, *C. difficile* spores were collected from 2-week Clospore media cultured bacteria and purified with sucrose gradient layer (50%, 45%, 35%, 25%, 10%). The purified spores were diluted to an OD_600_ of 1.0 in the germination buffer [10 mM Tris (pH 7.5), 150 mM NaCl, 100 mM glycine, 10 mM taurocholic acid (TA)] to detect the germination ratio. The value of OD_600_ was monitored immediately (0 min, t_0_) and detected once every 2 min (t_x_) for 20 min at 37 □. The germination ratio was calculated as OD_600_ (tx) / OD_600_ (T_0_). Spores in germination buffer without TA were used as a negative control. To evaluate the transcription of regulation genes during sporulation and germination, expression of *spo0A*, *sigE*, *sigF*, and *sigG* in sporulation process and *cspBA*, *cspC*, and *sleC* in spores germination process were detected by RT-qPCR.

*C. difficile* vegetative cell length at 6 and 10 h was detected by phase-contrast microscopy as described above. At least three fields of view for each strain were acquired with a DS-Fi2 camera from three independent experiments. The cell length was calculated by the imagining system software. To visualize cell separation, Fm4-64 Dye was added to *C. difficile* to a final concentration of 5 μg/ml and incubated on ice for 1 minute. Slides were prepared by placing 2 μl of *C. difficile* sample on a thin 1% agarose pad placed directly on top of the slide. Fluorescent microscopy was performed using an Olympus BX53 digital upright microscope and images were acquired with an Olympus DP26 color camera.

### Evaluation of virulence of R20291 and R20291Δ0971 in the mouse model of *C. difficile* infection

C57BL/6 female mice (6 weeks old) were ordered from Charles River Laboratories, Cambridge, MA. All studies were approved by the Institutional Animal Care and Use Committee of University of South Florida. The experimental design and antibiotic administration were conducted as previously described (Sun et al., 2011). Briefly, 20 mice were divided into 2 groups in 4 cages. Group 1 mice were challenged with R20291 spores and group 2 mice with R20291Δ0971 spores, respectively. Mice were given an orally administered antibiotic cocktail (kanamycin 0.4 mg ml^−1^, gentamicin 0.035 mg ml^−1^, colistin 0.042 mg ml^−1^, metronidazole 0.215 mg ml^−1^, and vancomycin 0.045 mg ml^−1^) in drinking water for 4 days. After 4 days of antibiotic treatment, all mice were given autoclaved water for 2 days, followed by one dose of clindamycin (10 mg kg^−1^, intraperitoneal route) 24 h before spores challenge (Day 0). After that, mice were orally gavaged with 10^6^ of spores and monitored daily for a week for changes in weight, diarrhea, mortality, and other symptoms of the disease.

### Determination of *C. difficile* spores and toxin levels in feces

Fecal pellets from post infection day 0 to day 7 were collected and stored at −80 □. To enumerate *C. difficile* numbers, feces were diluted with PBS at a final concentration of 0.1 g ml^−1^, followed by adding 900 μl of absolute ethanol into 100 μl of the fecal solution, and kept at room temperature for 1 h to inactivate vegetative cells. Afterward, fecal samples were serially diluted and plated on BHIS-CCT plates (250 μg ml^−1^ D-cycloserine, 8 μg ml^−1^ cefoxitin, 0.1% TA). After 48 h incubation, colonies were counted and expressed as CFU/g feces. To evaluate toxin tilter in feces, 0.1 g ml^−1^ of the fecal solution was diluted two times with PBS, followed by examining TcdA and TcdB ELISA test.

### Statistical analysis

The reported experiments were conducted in independent biological triplicates with the exception of animal experiments, and each sample was additionally taken in technical triplicates. Animal survivals were analyzed by Kaplan-Meier survival analysis and compared by the Log-Rank test. Student’s unpaired *t*-test was used for two groups comparison. One-way analysis of variance (ANOVA) was used for more than two groups comparison. Results were expressed as mean ± standard error of the mean. Differences were considered statistically significant if *P* < 0.05 (*).

## Acknowledgments

This work was supported in part by the National Institutes of Health grants (R01-AI132711, R01-AI149852, R01AI081062, and R01-AI124458).

## Supplement files

**Fig. S1. Visualization of Triton X-100 induced *C. difficile* autolysis with phase-contrast microscopy and DAPI / PI staining.**

*C. difficile* cells were grown for 12h. 1 ml of each culture was collected and centrifuged. The pellets were resuspended in a final volume of 100 μl of 50 mM potassium phosphate buffer (pH 7.0) with or without 0.1% of Triton X-100 and incubated at room temperature for 30 min. Then, the cells were visualized by phase-contrast microscopy and stained with DAPI / PI, respectively. A panel: R20291; B panel: R20291Δ0971; C panel: R20291Δ0971-C.

**Fig. S2. Detection of cell wall and septa by TEM**

Ultrathin 90 nm sectioned of *C. difficile* vegetative cell samples were stained with uranyl acetate and were examined with a JOEL JEM1400 transmission electron microscope using an accelerating voltage of 80 kV at 50 k×, 120 k×, and 200 k× magnification.

**Fig. S3. Visualization of spores and detection of sporulation ratio.**

Spores were detected by phase-contrast microscopy. At least three fields of view for each strain were acquired with a DS-Fi2 camera and used to calculate the percentage of spores (the number of spores divided by the total number of spores, prespores, and vegetative cells) from three independent experiments. A panel: R20291; B panel: R20291Δ0971; C panel: R20291Δ0971-C.

**Fig. S4. Cell length and separation detected by phase-contrast microscopy.**

*C. difficile* cultures from 6 and 10 h post-inoculation were used for slide preparation. At least three fields of view for each strain were acquired with a DS-Fi2 camera from three independent experiments. The cell length was calculated by imagining system software. 408 of R20291, 399 of R20291Δ0971, 402 of R20291Δ0971-C cells were counted, respectively. Phase-contrast microscopy (A and B), A panel: 6 h sample; B panel: 10 h sample. Fluorescent Phase-contrast microscopy (C and D), C panel: 6 h sample, D: 10 h sample.

**Fig. S5. Biofilm, adhesin, spore resistance to heat, and spore structure assay**

A. Biofilm formation assay. Biofilm formation of *C. difficile* strains was detected at 24 and 48 h, respectively.

B. Adhesion assay. The adhesion ability of *C. difficile* vegetative cells was determined on HCT-8 cells.

C. Spores resistance to heat. 1×10^6^ spores were used to detect spore resistance at 65 □ for 4 h.

D. Spore structure detected by TEM at 150 and 200 k× magnification.

Experiments were independently repeated thrice. Bars stand for mean ± SEM. One-way analysis of variance (ANOVA) with post-hoc Tukey test was used for statistical significance, statistically significant outcomes are indicted (\**P* < 0.05, \*\**P* < 0.01, \*\*\**P* < 0.001, \*\*\*\**P* < 0.0001).

**Fig. S6** *C. difficile* **motility assay on BHIS plates.**

*C. difficile* strains were cultured to an OD_600_ of 0.8. Then 2 μl of each cultures were penetrated into soft BHIS agar (0.175%) plate for swimming analysis, and 2 μl of each cultures were dropped onto 0.3% BHIS agar plate for swarming analysis. A. Swarming analysis at 24 h with 0.3% agar BHIS plate. 1: R20291; 2: R20291Δ0971; 3: R20291Δ0971-C. B. Swimming analysis at 12 h with 0.175% agar BHIS plate. 1: R20291; 2: R20291Δ0971; 3: R20291Δ0971-C. C. Swarming analysis at 24 h with 0.3% agar BHIS-Tm plate. 1: R20291-PMTL84153; 2: R20291Δ0971-pMTL84153; 3: R20291Δ0971-C. D. Swimming analysis at 12 h with 0.175% agar BHIS-Tm plate. 1: R20291-PMTL84153; 2: R20291Δ0971-pMTL84153; 3: R20291Δ0971-C. E. Diameter of motility zone.

Experiments were independently repeated thrice. Bars stand for mean ± SEM. One-way analysis of variance (ANOVA) with post-hoc Tukey test was used for statistical significance, statistically significant outcomes are indicted (\**P* < 0.05, \*\**P* < 0.01, \*\*\**P* < 0.001).

**Table S1.**
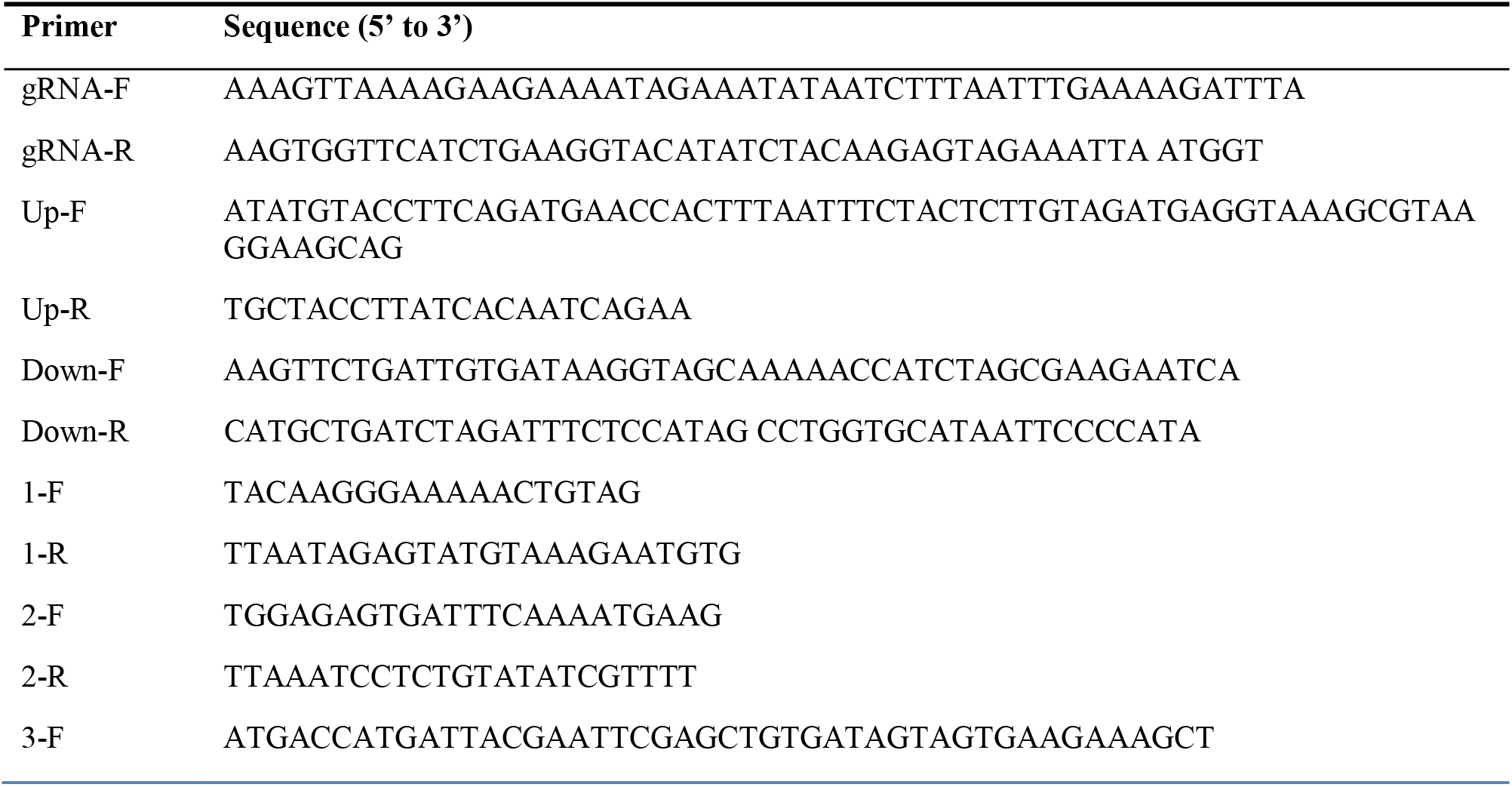
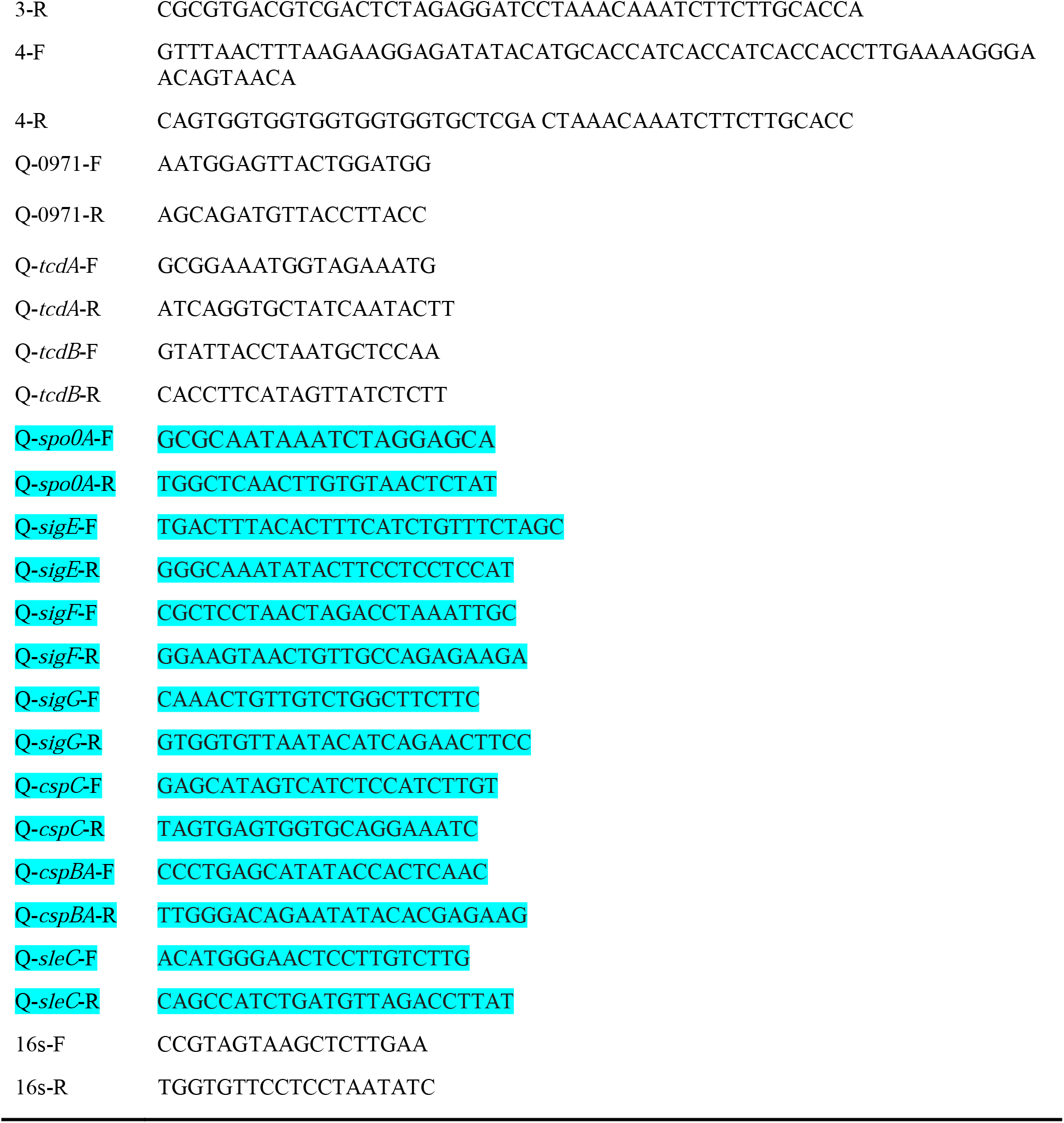
Primers utilized in this study.

