## Supplemental figures and tables for "Cwl0971, a novel peptidoglycan hydrolase, plays pleiotropic roles in *Clostridioides difficile* R20291"

**Supplement files**

**Support methods**

**Biofilm assay**

For biofilm analysis, *C. difficile* strains were cultured to an OD_600_ of 0.8, and 1% of *C. difficile* cultures were inoculated into Reinforced Clostridial Medium (RCM) with 8 well repeats in 96-well plate and incubated in the anaerobic chamber at 37 ℃ for 24 and 48 h. The formation of biofilm was analyzed by crystal violet dye. Briefly, *C. difficile* cultures were removed by pipette carefully. Then 100 µl of 2.5% glutaraldehyde was added into the well to fix the bottom biofilm, and the plate was kept at room temperature for 30 min. Next, the wells were washed 3 times with PBS and dyed with 0.25% (w/v) crystal violet for 10 min. The crystal violet solution was removed, and the wells were washed 5 times with PBS, followed by the addition of acetone into wells to dissolve the crystal violet of the cells. The dissolved solution was further diluted with ethanol 2 - 4 times and then detected at OD_570_.

**Adhesion of *C. difficile* vegetative cells to HCT-8 cells**

*C. difficile* adhesion ability was evaluated with HCT-8 cells (ATCC CCL-244) (Janvilisri et al., 2010). Briefly, HCT-8 cells were grown to 95% confluence (2 × 10^5^/well) in a 24-well plate and then moved into the anaerobic chamber, followed by infecting with 6 × 10^6^ of log phase of *C. difficile* vegetative cells at a multiplicity of infection (MOI) of 30:1. The plate was cultured at 37 ℃ for 30 min. After incubation, the infected cells were washed with 300 µl of PBS 3 times, and then suspended in RPMI media with trypsin and plated on BHIS agar plates to enumerate the adhered *C. difficile* cells. The adhesion ability of *C. difficile* to HCT-8 cells was calculated as follow: CFU of adhered bacteria / total cell numbers.

**Spore resistance to heat**

To evaluate the spore resistance to heat, the sucrose gradient-purified *C. difficile* spores were shocked in a 65 ℃ water bath for 1 - 4 h. Afterward, the spores were plated on BHIS plates with 0.1% TA. The survival rate was calculated as CFU (65 ℃ heated) / CFU (no heated).

**Motility assay**

To examine the effect of 0971 gene deletion on *C. difficile* motility, *C. difficile* strains were cultured to an OD_600_ of 0.8. For swimming analysis, 2 µl of *C. difficile* culture was penetrated into soft BHIS agar (0.175%) plates, meanwhile, 2 µl of culture was dropped onto 0.3% BHIS agar plates for swarming analysis. The swimming assay plates were incubated for 12 h and the swarming plates were incubated for 24 h, respectively.

**Support figures and Table**

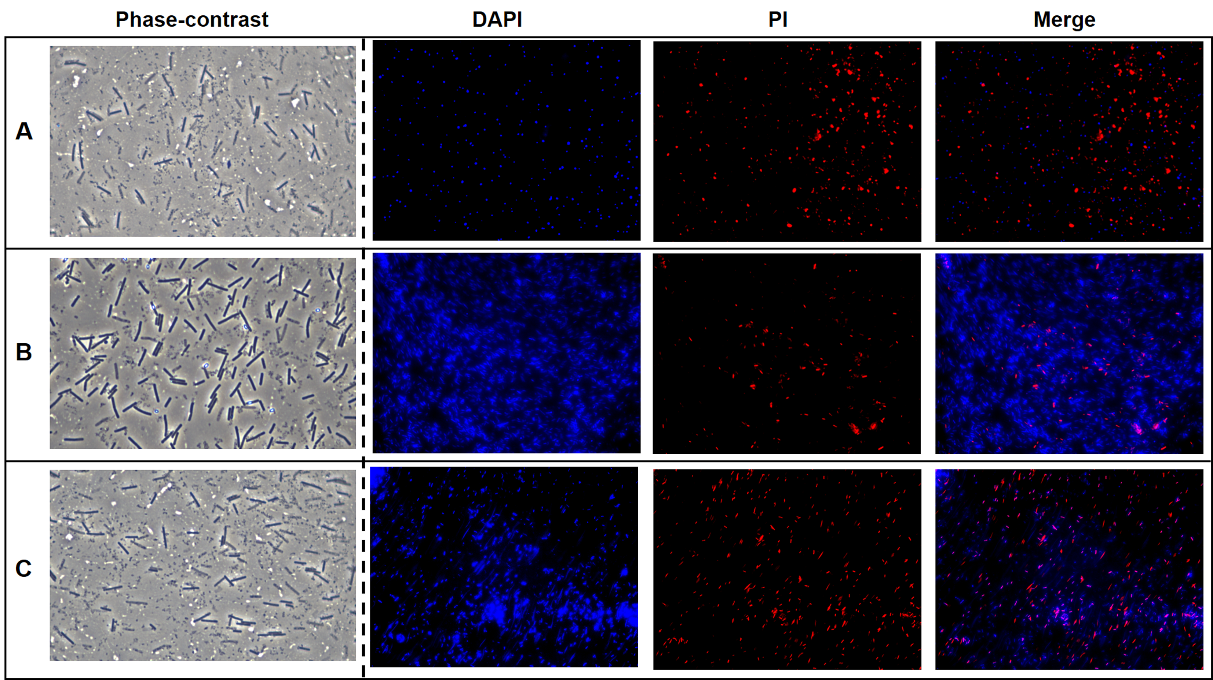

**Fig. S1. Visualization of Triton X-100 induced *C. difficile* autolysis with phase-contrast microscopy and DAPI / PI staining.**

*C. difficile* cells were grown for 12h. 1 ml of each culture was collected and centrifuged. The pellets were resuspended in a final volume of 100 µl of 50 mM potassium phosphate buffer (pH 7.0) with or without 0.1% of Triton X-100 and incubated at room temperature for 30 min. Then, the cells were visualized by phase-contrast microscopy and stained with DAPI / PI, respectively. A panel: R20291; B panel: R20291Δ0971; C panel: R20291Δ0971-C.

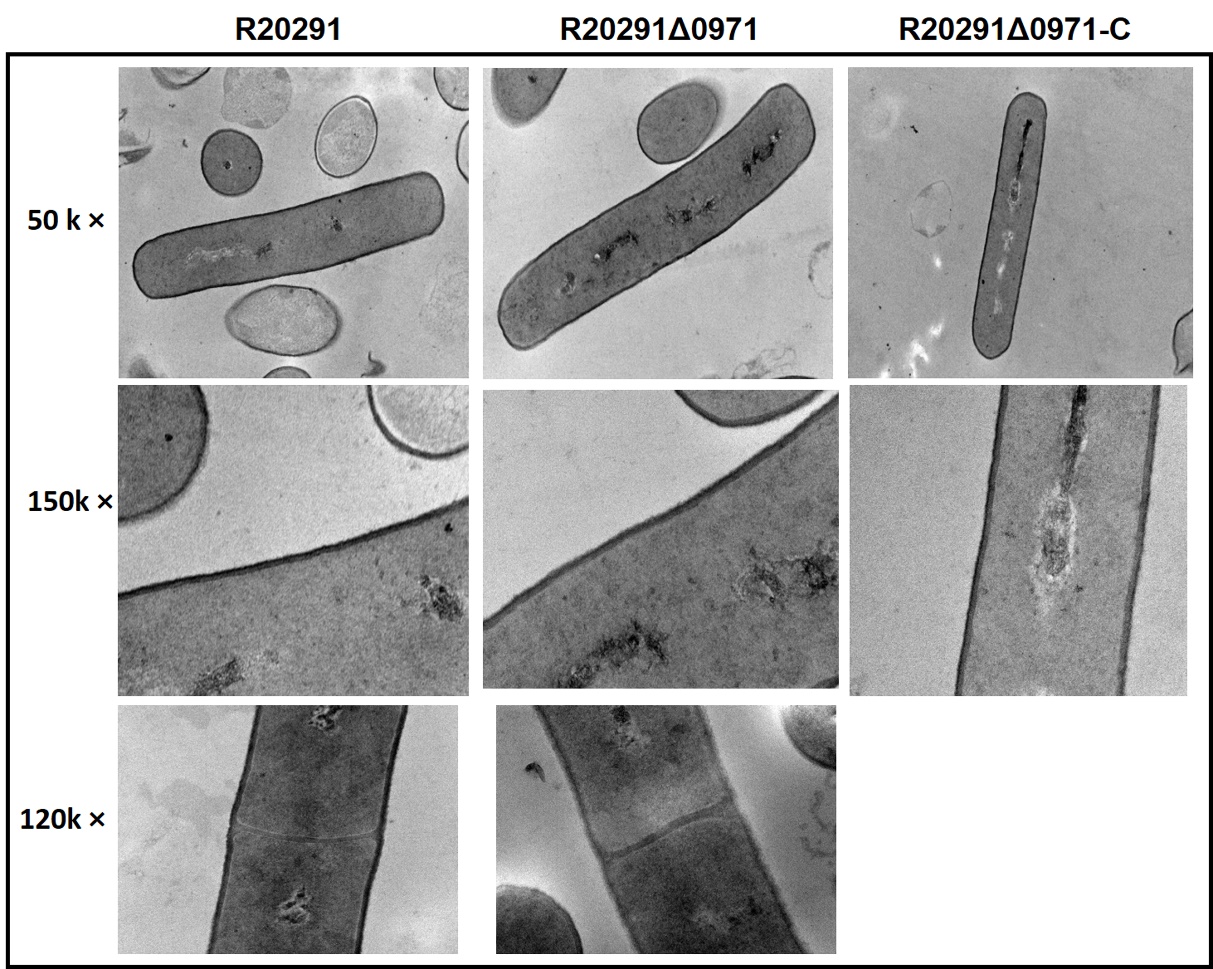

**Fig. S2.** **Detection of cell wall and septa by TEM**

Ultrathin 90 nm sectioned of *C. difficile* vegetative cell samples were stained with uranyl acetate and were examined with a JOEL JEM1400 transmission electron microscope using an accelerating voltage of 80 kV at 50 k×, 120 k×, and 200 k× magnification.

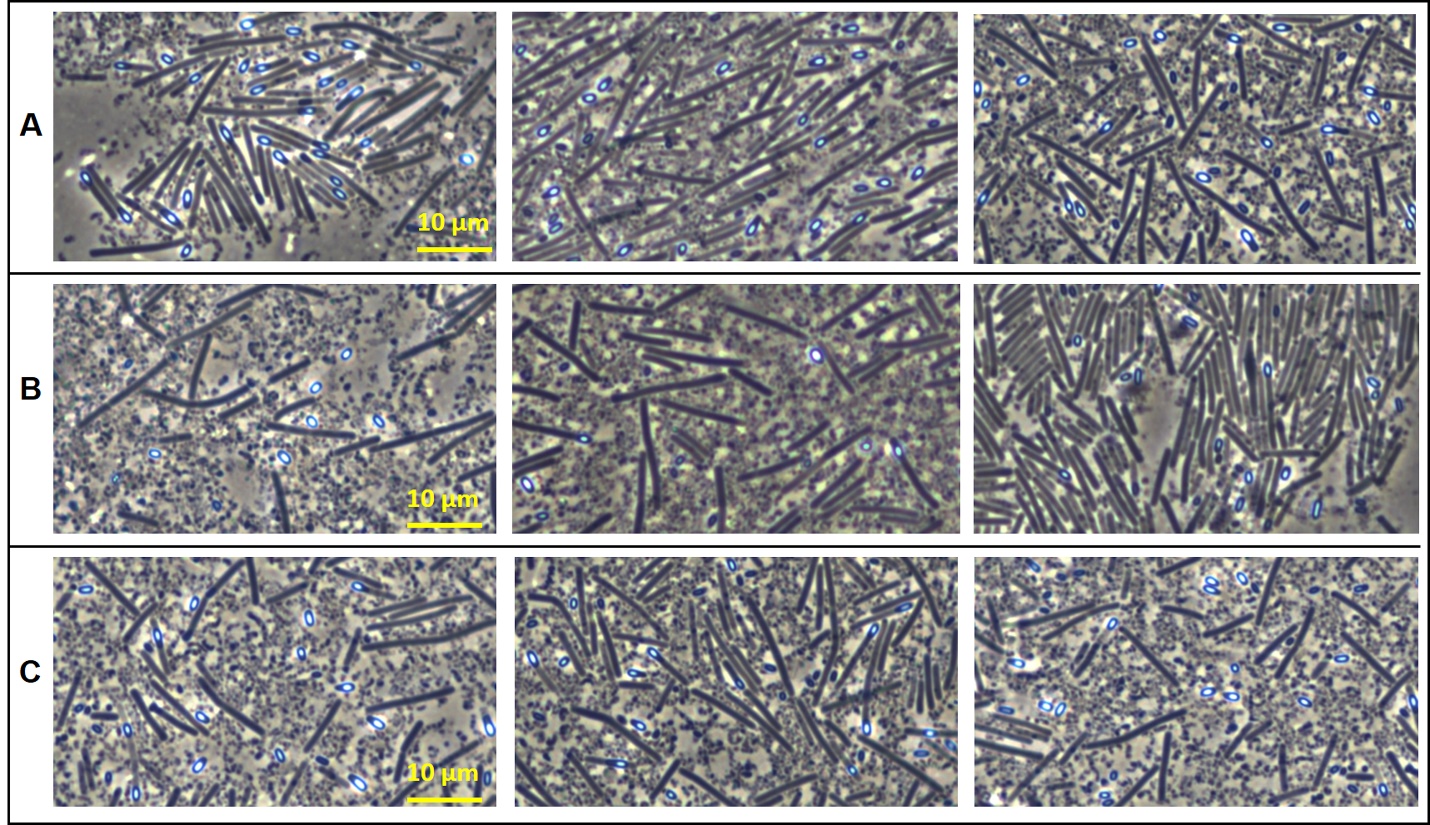

**Fig. S3.** **Visualization of spores and detection of sporulation ratio.**

Spores were detected by phase-contrast microscopy. At least three ﬁelds of view for each strain were acquired with a DS-Fi2 camera and used to calculate the percentage of spores (the number of spores divided by the total number of spores, prespores, and vegetative cells) from three independent experiments. A panel: R20291; B panel: R20291Δ0971; C panel: R20291Δ0971-C.

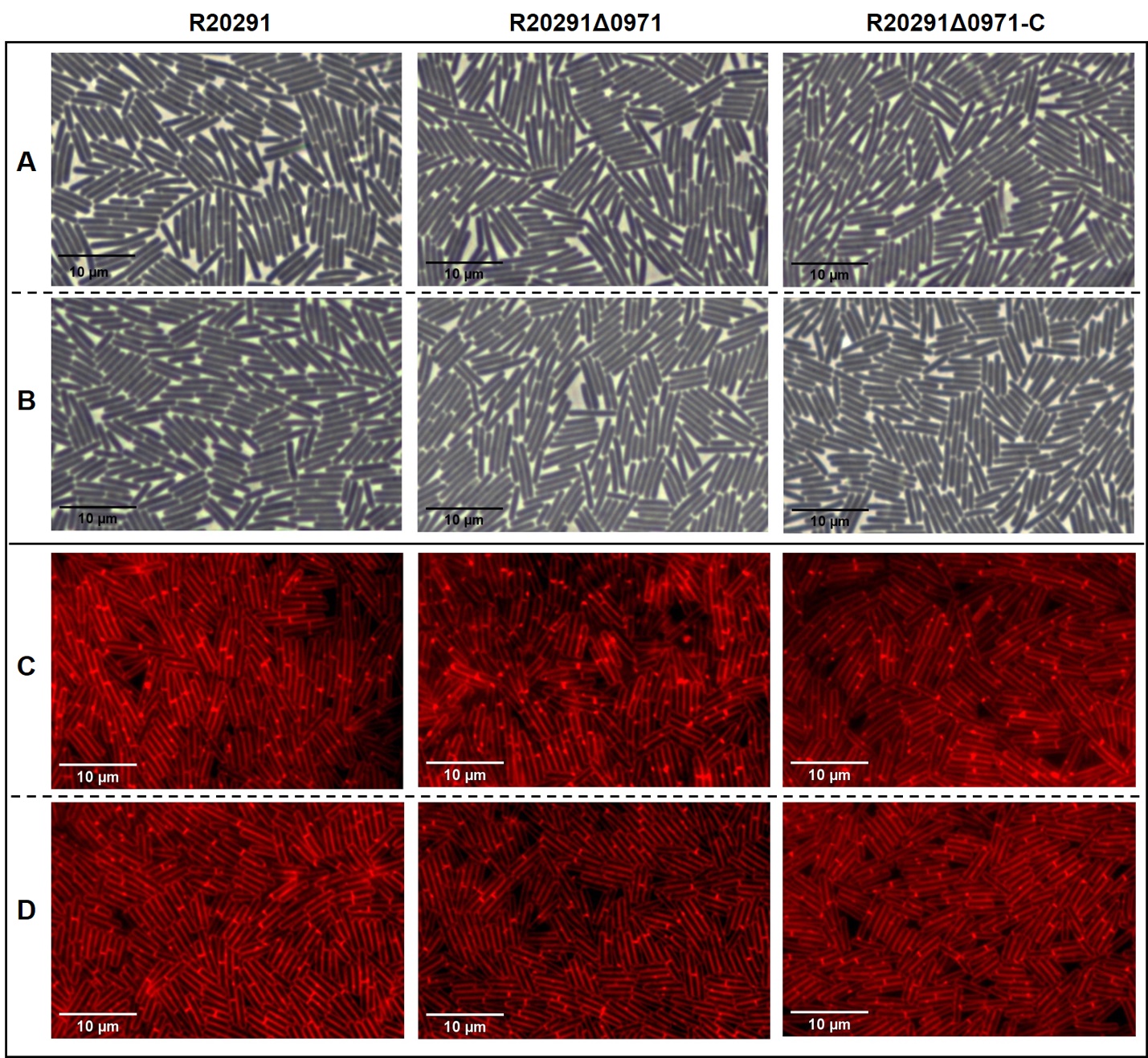

**Fig. S4. Cell length and separation detected by phase-contrast microscopy.**

*C. difficile* cultures from 6 and 10 h post-inoculation were used for slide preparation. At least three ﬁelds of view for each strain were acquired with a DS-Fi2 camera from three independent experiments. The cell length was calculated by imagining system software. 408 of R20291, 399 of R20291Δ0971, 402 of R20291Δ0971-C cells were counted, respectively. Phase-contrast microscopy (A and B), A panel: 6 h sample; B panel: 10 h sample. Fluorescent Phase-contrast microscopy (C and D), C panel: 6 h sample, D: 10 h sample.

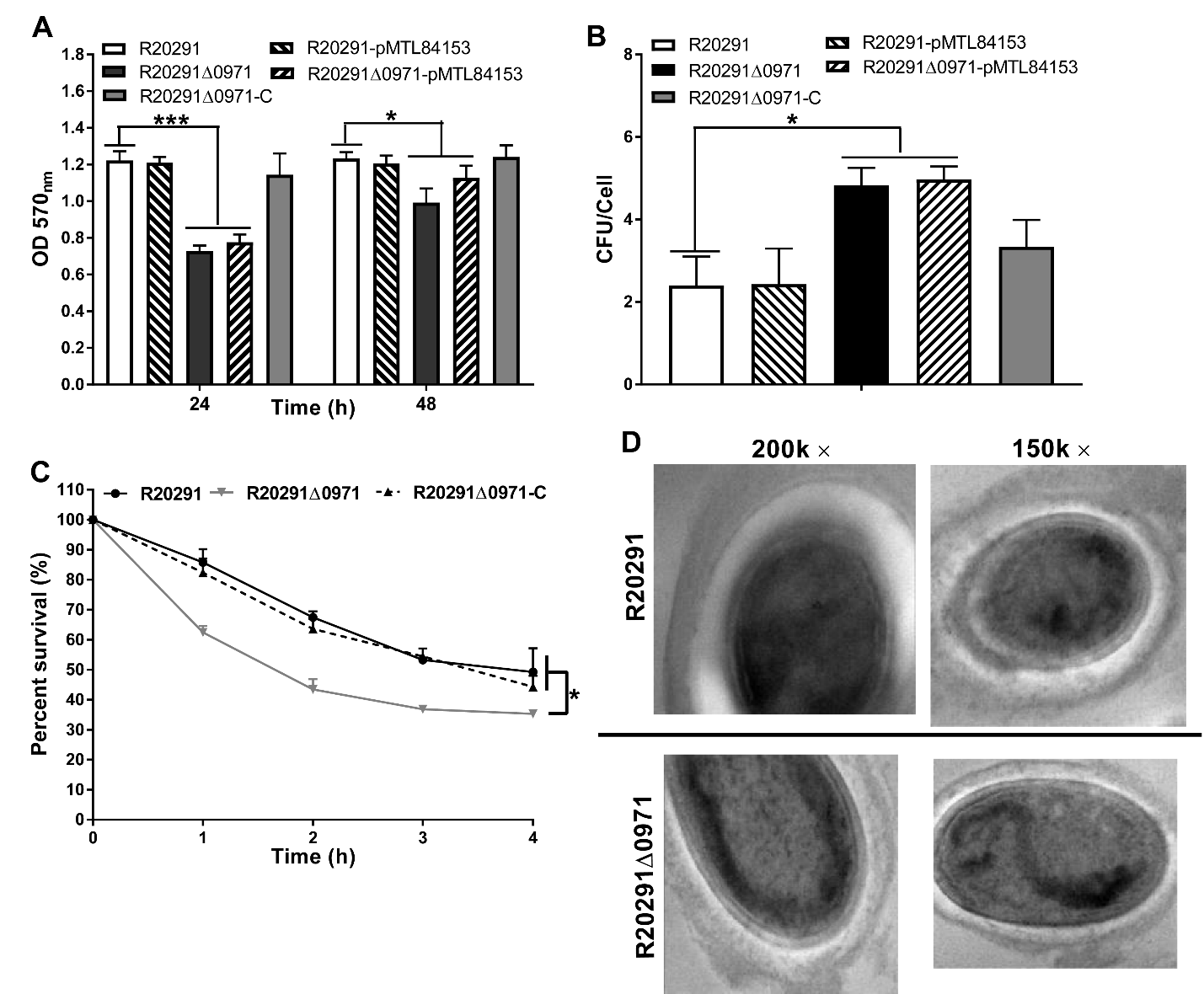

**Fig. S5. Biofilm, adhesin, spore resistance to heat, and spore structure assay**

A. Biofilm formation assay. Biofilm formation of *C. difficile* strains was detected at 24 and 48 h, respectively.

B. Adhesion assay. The adhesion ability of *C. difficile* vegetative cells was determined on HCT-8 cells.

C. Spores resistance to heat. 1×10^6^ spores were used to detect spore resistance at 65 ℃ for 4 h.

D. Spore structure detected by TEM at 150 and 200 k× magnification.

Experiments were independently repeated thrice. Bars stand for mean ± SEM. One-way analysis of variance (ANOVA) with post-hoc Tukey test was used for statistical significance, statistically significant outcomes are indicted (**P* < 0.05, ***P* < 0.01, ****P* < 0.001, *****P* < 0.0001).

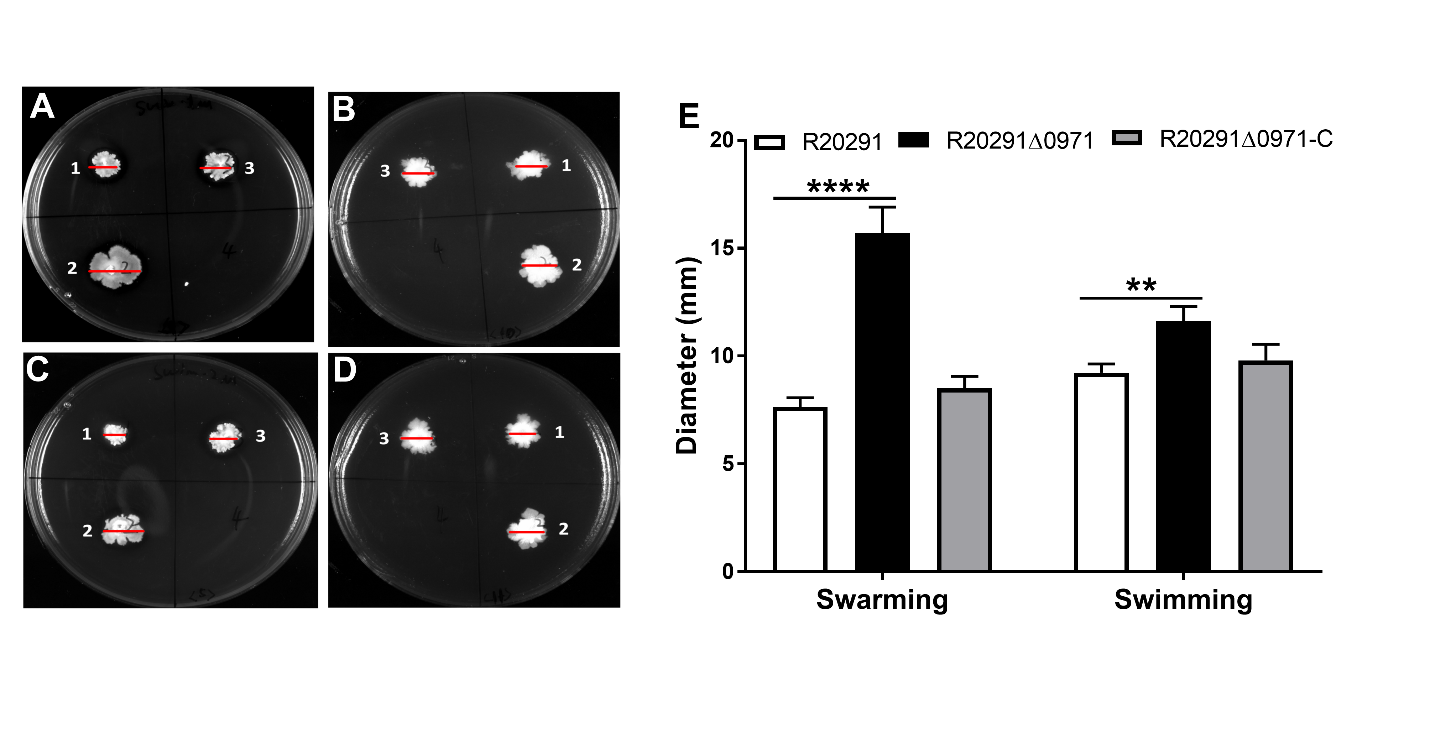

**Fig. S6 *C. difficile* motility assay on BHIS plates.**

*C. difficile* strains were cultured to an OD_600_ of 0.8. Then 2 µl of each cultures were penetrated into soft BHIS agar (0.175%) plate for swimming analysis, and 2 µl of each cultures were dropped onto 0.3% BHIS agar plate for swarming analysis. A. Swarming analysis at 24 h with 0.3% agar BHIS plate. 1: R20291; 2: R20291Δ0971; 3: R20291Δ0971-C. B. Swimming analysis at 12 h with 0.175% agar BHIS plate. 1: R20291; 2: R20291Δ0971; 3: R20291Δ0971-C. C. Swarming analysis at 24 h with 0.3% agar BHIS-Tm plate. 1: R20291-PMTL84153; 2: R20291Δ0971-pMTL84153; 3: R20291Δ0971-C. D. Swimming analysis at 12 h with 0.175% agar BHIS-Tm plate. 1: R20291-PMTL84153; 2: R20291Δ0971-pMTL84153; 3: R20291Δ0971-C. E. Diameter of motility zone.

Experiments were independently repeated thrice. Bars stand for mean ± SEM. One-way analysis of variance (ANOVA) with post-hoc Tukey test was used for statistical significance, statistically significant outcomes are indicted (**P* < 0.05, ***P* < 0.01, ****P* < 0.001).

**Table S1. Primers utilized in this study.**

| **Primer** | **Sequence (5’ to 3’)** |
| --- | --- |
| gRNA-F | AAAGTTAAAAGAAGAAAATAGAAATATAATCTTTAATTTGAAAAGATTTA |
| gRNA-R | AAGTGGTTCATCTGAAGGTACATATCTACAAGAGTAGAAATTA ATGGT |
| Up-F | ATATGTACCTTCAGATGAACCACTTTAATTTCTACTCTTGTAGATGAGGTAAAGCGTAAGGAAGCAG |
| Up-R | TGCTACCTTATCACAATCAGAA |
| Down-F | AAGTTCTGATTGTGATAAGGTAGCAAAAACCATCTAGCGAAGAATCA |
| Down-R | CATGCTGATCTAGATTTCTCCATAG CCTGGTGCATAATTCCCCATA |
| 1-F | TACAAGGGAAAAACTGTAG |
| 1-R | TTAATAGAGTATGTAAAGAATGTG |
| 2-F | TGGAGAGTGATTTCAAAATGAAG |
| 2-R | TTAAATCCTCTGTATATCGTTTT |
| 3-F | ATGACCATGATTACGAATTCGAGCTGTGATAGTAGTGAAGAAAGCT |
| 3-R | CGCGTGACGTCGACTCTAGAGGATCCTAAACAAATCTTCTTGCACCA |
| 4-F | GTTTAACTTTAAGAAGGAGATATACATGCACCATCACCATCACCACCTTGAAAAGGGAACAGTAACA |
| 4-R | CAGTGGTGGTGGTGGTGGTGCTCGA CTAAACAAATCTTCTTGCACC |
| Q-0971-F | AATGGAGTTACTGGATGG |
| Q-0971-R | AGCAGATGTTACCTTACC |
| Q-*tcdA*-F | GCGGAAATGGTAGAAATG |
| Q-*tcdA*-R | ATCAGGTGCTATCAATACTT |
| Q-*tcdB*-F | GTATTACCTAATGCTCCAA |
| Q-*tcdB*-R | CACCTTCATAGTTATCTCTT |
| Q-*spo0A*-F | GCGCAATAAATCTAGGAGCA |
| Q-*spo0A*-R | TGGCTCAACTTGTGTAACTCTAT |
| Q-*sigE*-F | TGACTTTACACTTTCATCTGTTTCTAGC |
| Q-*sigE*-R | GGGCAAATATACTTCCTCCTCCAT |
| Q-*sigF*-F | CGCTCCTAACTAGACCTAAATTGC |
| Q-*sigF*-R | GGAAGTAACTGTTGCCAGAGAAGA |
| Q-*sigG*-F | CAAACTGTTGTCTGGCTTCTTC |
| Q-*sigG*-R | GTGGTGTTAATACATCAGAACTTCC |
| Q-*cspC*-F | GAGCATAGTCATCTCCATCTTGT |
| Q-*cspC*-R | TAGTGAGTGGTGCAGGAAATC |
| Q-*cspBA*-F | CCCTGAGCATATACCACTCAAC |
| Q-*cspBA*-R | TTGGGACAGAATATACACGAGAAG |
| Q-*sleC*-F | ACATGGGAACTCCTTGTCTTG |
| Q-*sleC*-R | CAGCCATCTGATGTTAGACCTTAT |
| 16s-F | CCGTAGTAAGCTCTTGAA |
| 16s-R | TGGTGTTCCTCCTAATATC |
